# A miRNA screen identifies a transcriptional program controlling the fate of adult stem cell

**DOI:** 10.1101/314658

**Authors:** Jacques Robert, Efstathios Vlachavas, Gilles Lemaître, Aristotelis Chatziioannou, Michel Puceat, Frederic Delom, Delphine Fessart

## Abstract

The 3D cultures provide more insight into cell-to-cell and cell-to-matrix interactions, better mimicking the environment where stem cells reside compared to traditional 2D cultures. Although the precise molecular pathways involved in the regulation of stem and progenitor cell fate remain unknown, it is widely accepted that transcription factors play a crucial role as intrinsic regulators in these fate decisions.

In this study, we carried out a microRNA screen to track the behaviour of adult stem/progenitor cells derived from human mammary epithelial cells grown in 3D cultures. We identified miR-106a-3p, which enriches the adult stem cell-like lineage and promotes the expansion of 3D cultures. Transcriptomic analysis showed that this miRNA regulates transcription factors such as REST, CBFB, NF-YA, and GATA3, thereby enhancing the maintenance of adult stem/progenitor cells in human epithelial cells. These data reveal a clear transcriptional program that governs the maintenance of adult stem/progenitor cells and controls their fate.

## Introduction

Three-dimensional (3D) human culture models are appealing tools for studying pathophysiological processes. Such models have been developed by our team and others for the lung [1, 2], as well as for various other organs [3]. These *in vitro* models offer an effective method to assess stem cell behaviour and investigate their molecular regulation. Adult stem cells (aSCs) exhibit a high degree of specialisation, allowing them to differentiate into multiple cell types. However, they can only generate tissues in which they reside, classifying them as either ‘unipotent’ or ’multipotent’. This classification highlights their limited versatility compared to pluripotent cells. Pluripotent cells, such as embryonic stem cells (ESCs), can give rise to all cell types. Consistent with this notion, primary cell populations enriched with known progenitor/stem cell markers are more efficient at forming 3D structures than general cell populations [4]. The development and enrichment of breast stem cells likely depend on the coordination of multiple critical transcriptional processes. Interestingly, increasing amounts of data show that the same molecular pathways regulate the self-renewal of both normal stem cells and Cancer Stem Cells (CSCs) in tumours [5]. CSCs express components of the core pluripotency complex found in ESCs, including octamer-binding transcription factor-4 (OCT4), SRY-Box Transcription Factor 2 (SOX2), and NANOG. These factors are closely associated with CSC development [6–11]. OCT4 expression has been reported in both differentiated normal and malignant human cells [12]. SOX2 expression is observed in various malignant tissues [13], and NANOG expression is detected in human neoplasms, including germ cell tumours, breast carcinoma, and osteosarcoma [14]. Additionally, SOX2 has been found in tumour-spheres derived from breast cancer tumours and cell lines [15]. These observations suggest that adult breast stem cells may contain cells with properties similar to embryonic-like stem cells [14, 16]. Currently, it is not known whether adult stem cells share a comparable molecular signature of stemness, such as a minimal core transcriptional program.

MicroRNAs (miRNAs) have been shown to play a crucial role in regulating stem cell self-renewal and differentiation [17]. Generally, a single gene can be repressed by multiple miRNAs, while one miRNA may target and repress multiple genes, which results in the formation of complex regulatory networks. In many developmental processes, miRNAs finely regulate cellular identities by targeting key transcription factors in critical pathways [18]. To investigate the contribution of miRNA-mediated gene regulation in maintaining the adult stem cell-like lineage, we aimed to identify the mechanisms controlling the cell- initiating subpopulation and optimise tissue-specific 3D growth conditions. Since 3D structures originate from individual stem/progenitor cells, we conducted a functional miRNA screen on human primary epithelial cells to identify factors critical to the formation and expansion of aSC-derived 3D structures. In this study, we identified miR-106a-3p and its target genes as key players in these processes. Transcriptomic profiling of miR-106a-3p– transduced cells revealed genetic programs overlapping with those found in other stem and progenitor cells, displaying common features with ESC-like cells or intermediate cellular states. Through a gain-of-function approach, we demonstrated that endogenous levels of three core transcription factors—OCT4, SOX2, and NANOG—are essential for generating 3D structures from primary cells. Furthermore, we identified the mechanism by which miR-106a- 3p finely tunes the differentiation process through a set of transcription regulators, including CBFB (Core-binding factor b), NFYA (Nuclear Transcription Factor Y Subunit Alpha), GATA3 (GATA Binding Protein 3), and REST (RE1 silencing transcription factor). These regulators are critical for maintaining the aSC-like lineage. In conclusion, our results highlight the significant role of miR-106a-3p in sustaining the aSC-derived 3D structure and elucidate the transcriptional mechanisms underlying their growth.

## Results

### 3D cultures derived from normal human mammary epithelial cells preserve some mammary epithelial lineages

The mammary epithelium is composed of distinct cell lineages among which luminal epithelial and myoepithelial cells. The choice of growth media and culture methods is critical in shaping the characteristics of primary human mammary epithelial cell strains and determining experimental outcomes. Finite-lifespan human mammary epithelial cells (HMECs) were generously provided by M. Stampfer via the Human Mammary Epithelial Cell (HMEC) Bank and were cultured in M87A-type medium. M87A medium has been noted for its ability to support pre-stasis HMEC growth for up to 60 population doublings (PD) and maintain luminal cells through approximately 8 passages, equivalent to about 30 PD [19]. To evaluate the heterogeneity of pre-stasis HMECs cultured in M87A, we performed immunostaining for epithelial lineage markers, as previously described [19]. We confirmed the presence of luminal epithelial progenitor (LEP) and myoepithelial progenitor (MEP) cells using specific antibodies against lineage-specific markers (CK18 and MUC1 for LEP cells, and CK14 and CDK5 for MEP cells) (Figure 1A). In addition, we evaluated aldehyde dehydrogenase (ALDH) activity as a marker for normal and malignant human mammary stem cells, as well as a predictor of clinical outcomes, in line with the findings of Ginestier *et al.*[20]. This analysis demonstrated that under these culture conditions, cells preserve stem/progenitor properties (Figure 1B). Using the ALDEFLUOR assay, we identified that approximately 5.2% of normal mammary epithelial cells exhibit ALDH enzymatic activity (Figure 1B). To further explore the regenerative potential of these cells, we performed single-cell seeding in a 3D matrix (Matrigel). This process led to the formation of small, initially spherical structures (up to Day 8) that later differentiated into organoids, with branching initiation observed by Day 10. By Day 20, the structures exhibited budding and lobule formation (Figure 1C), resembling the functional Terminal Duct Lobular Unit (TDLU) found in human breast tissue (Figure 1C). Under these 3D culture conditions, HMECs retained self-renewal capabilities and the ability to regenerate secondary structures over three generations (Figure 1D).

**Figure 1:**
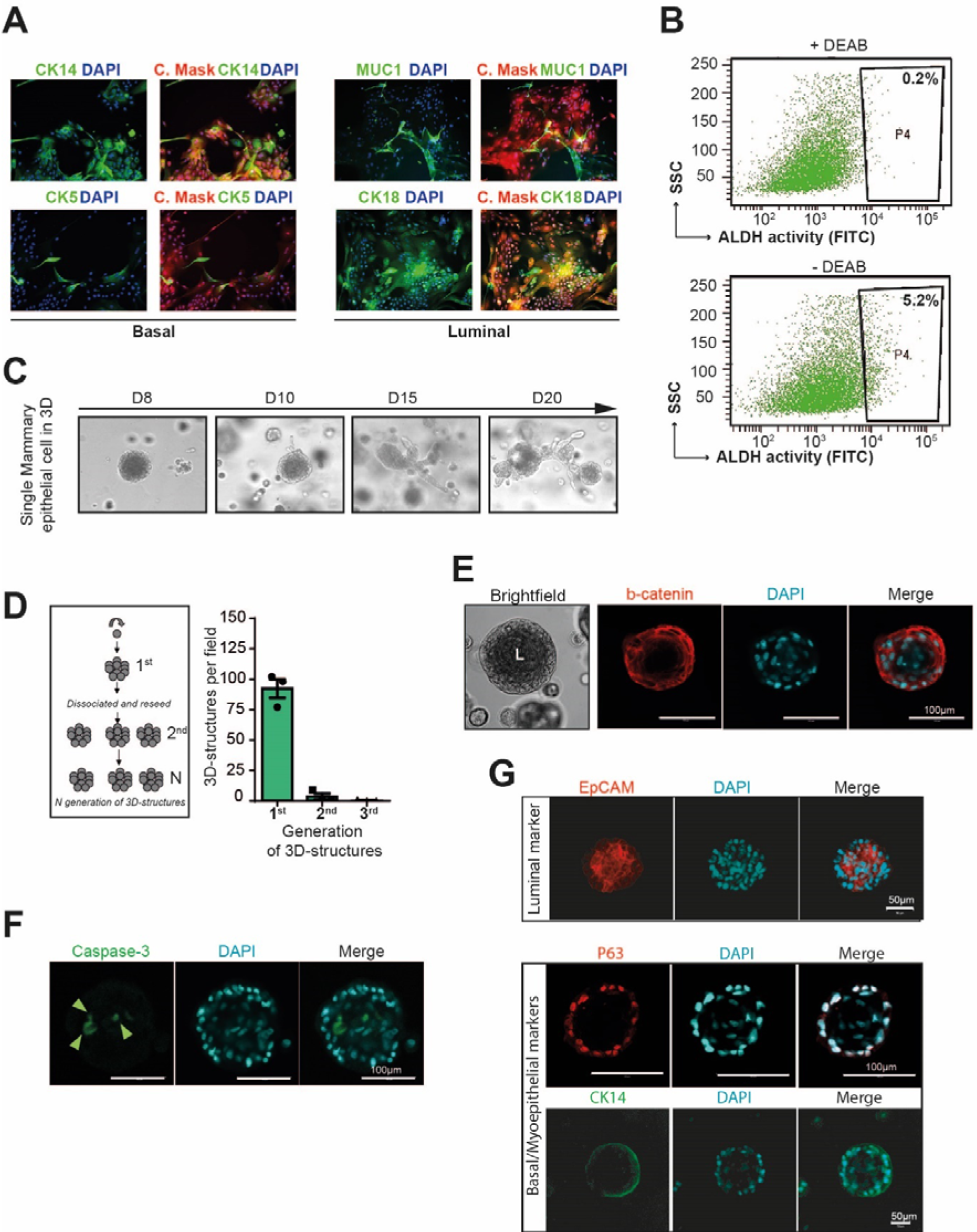
3D cultures of mammary primary epithelial cells. **A,** Representative immunofluorescence (IF) analysis depicting basal/myoepithelial markers stained for CK14 (green), CK5 (in green), entire cell with Cell Mask (red), and nuclei with DAPI (blue). Additionally, luminal markers stained for CK18 (green), MUC1 (green), cell mask (red), and DAPI. **B,** Representative Fluorescence-Activated Cell Sorting (FACS) analysis of normal breast epithelial cells using the ALDEFLUOR assay. Cells incubated with ALDEFLUOR substrate (BAAA) and the specific ALDH inhibitor, DEAB, established the baseline fluorescence and defined the ALDEFLUOR-positive region (P4) (n=3, mean ± SEM). **C,** Development of primary breast cells (single MECs) in 3D Matrigel following 20 days. Results are representative of at least three independent experiments. **D,** Quantification of primary, secondary, and 3rd generation 3D structure development. Data are presented as mean values ± SD, n=3. **E,** Immuno-fluorescent staining of β-catenin (red) and DAPI nuclear staining (blue) in 3D Matrigel organoids. Results are representative of three independent repeats of this experiment. **F,** Confocal cross-sections of 3D structure stained with active Caspase-3 and DAPI (blue) for the nucleus. Scale bars, 50 μm. **G,** Confocal cross-sections of 3D structures stained with Epcam (red) as the luminal marker and p63 (red), CK14 (green) as basal/myoepithelial markers, and DAPI (blue) for the nucleus. Scale bars, 50 μm.

Our next objective was to verify whether HMECs cultured in 3D Matrigel maintained key mammary structural characteristics. Confocal microscopy revealed that cells self-organised, forming a lumen within the 3D structures (Figure 1E). Tracking individual structures showed progressive lumen formation, reaching maximum size by Day 10 of culture (Figure 1F). Confocal sectioning of immuno-stained 3D structures for the luminal progenitor/mature luminal cell marker EpCAM and the basal/myoepithelial markers cytokeratin 14 (CK14) and p63 revealed that these structures were composed of luminal cells, basal cells, or a mixture of both (Figure 1G). This finding indicates that both basal and luminal cells are maintained in these cultures. Thus, primary cells cultured under these conditions preserve mammary epithelial lineages and retain the expression patterns of key mammary markers.

### 3D cultures derived from human mammary epithelial cells exhibit a CD44^high^/CD24^low^ phenotype

This study aimed to identify epithelial cell subpopulations capable of forming 3D structures and specifying stem/progenitor cell functions [4]. In this context, stem/progenitor cells are rare, immortal cells within a population of cultured cells. They possess the ability to both self-renew and generate other cell types through asymmetric cell division. Previous research has demonstrated that HMECs represent a multipotent stem cell population located in the basal layer of the mammary gland [21–23]. These cells are estrogen-independent (tamoxifen-resistant) and display heterogeneous expression of both luminal and myoepithelial lineage markers (Figure 1A) [24]. Initially, we compared the properties of HMECs cultured in 3D environments to those grown in conventional 2D culture systems. Approximately 3% of the cells in culture demonstrated the capacity to generate 3D structures (Figure 2A). The self-renewal capacity of 3D structure-initiating cells was then evaluated through serial passages, reflecting the maintenance of adult stem/progenitor cells from passage 5 to passage 11 (Figure 2A). As cells were serially propagated, their ability to form 3D structures progressively diminished, consistent with the previously observed loss of self-renewal potential in primary epithelial stem/progenitor cells after a few passages [25].

**Figure 2.**
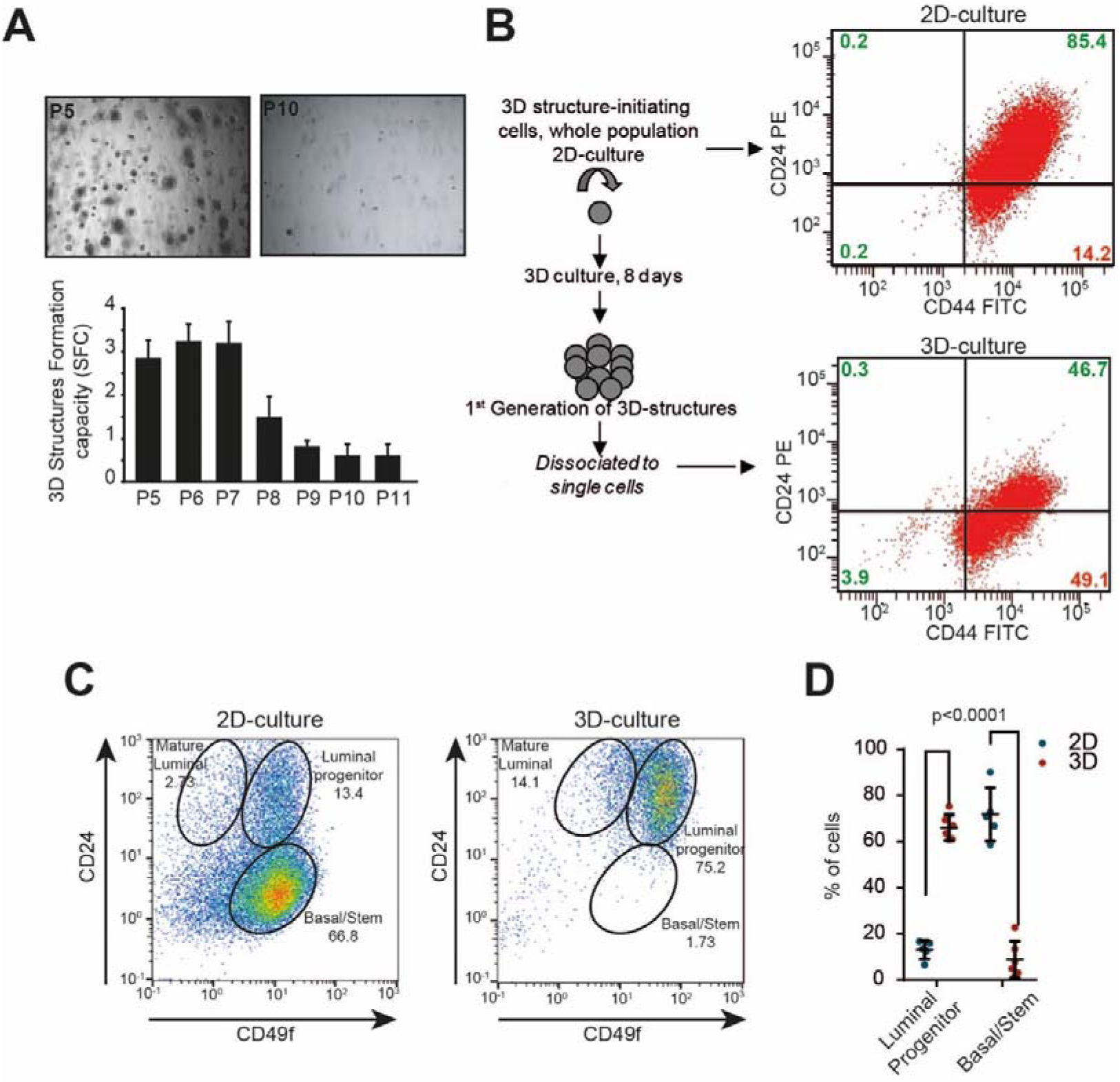
3D-culture confers original cell properties as compared to 2D-culture. **A**, Representative brightfield pictures of primary cells grown in 3D at passage 5 (P5) and passage 10 (P10). The bar graph shows the SFC, the mean ± SEM of 3D structure per well, from passage P5 to passage P11. Data are from three independent experiments for each passage. **B**, Flow cytometry analyses of CD44/CD24 in HMEC cells derived from 2D-cell culture (top) or from primary culture in 3D (bottom). The expression of CD44^high^/CD24^low^ in dissociated 3D structures was higher than in 2D cultured cells. A minimum of 10,000 events were collected per sample. (n=3, mean ± SEM) **C**, Representative flow cytometry analysis of CD24/CD49f in HMEC culture in 2D (left) compared to 3D (right). Mature luminal, luminal progenitor, and basal/stem populations are indicated. (n=3, mean ± SEM) **D**, Mammary epithelial cell population changes between cell culture conditions. Percent of Luminal progenitor (C24^high^/CD49f^high^) and basal/stem cells (CD24^low^/CD49f^high^) are quantified between culture conditions. Data are from four to five independent experiments. (mean±SEM).

Previous studies have highlighted that HMECs with a CD44^high^/CD24^low^ phenotype possess the highest progenitor activity compared to other stem/progenitor subpopulations [26]. This small subpopulation, characterized by self-renewal and a high proliferation rate, originates from normal stem cells and drives the formation of 3D structures [27–30]. Consequently, the CD44^high^/CD24^low^ phenotype has been widely used in combination with other markers as reliable marker for isolating normal breast adult stem cells [31–34]. To confirm the generation of stem-like cells in 3D culture, we used flow cytometry to assess the expression of breast stem cell markers (CD44^high^/CD24^low^) in 3D-grown HMECs, comparing them to cells grown in 2D cultures (Figure 2B). In 2D cultures, 85% of cells expressed high levels of both CD24 and CD44 (CD44^high^/CD24^high^), while 14% displayed the CD44^high^/CD24^low^ phenotype (Figure 2B, top panel). By contrast, 3D cultures showed a more than threefold increase in CD44^high^/CD24^low^ cells (∼49%) (Figure 1C, lower panel; p = 0.0268, n = 3). To further compare lineage specification between 2D and 3D culture conditions, we analysed pre-stasis HMECs by FACS for CD24^low^ and CD49f expression. CD49f, in combination with CD44^high^/CD24^low^, serves as a marker to segregate different cell lineages. In 3D cultures, we observed an enrichment of luminal progenitors, while in 2D cultures, basal/stem cells were more prevalent (Figure 2C). Specifically, 2D cultures exhibited significantly more basal/stem cells and less luminal progenitors, whereas 3D cultures displayed the opposite trend, with more luminal progenitors and less basal/stem cells (Figure 2D). Together, these findings suggest that cells grown under 3D culture conditions acquire a CD44^high^/CD24^low^ expression pattern similar to that of stem/progenitor cells. This indicates that 3D culture can be a valuable tool for enriching luminal stem/progenitor cell markers, facilitating further screening and research.

### A miRNA screening approach to selectively enhance the generation of 3D structure

To investigate whether miRNA-mediated gene regulation could enhance the ability to form 3D structures, we designed a two-step functional screening strategy aimed at enriching the CD44^high^/CD24^low^ cell population and increasing the number of 3D structure-generating cells. We monitored the expression of CD44 and CD24 after transfecting HMEC cells with miRNAs (Figure 3A-D). After performing quantitative image analysis on over 100,000 cells at Passage 6 (P6), the frequency distribution of CD44 intensity was compared between mass-cultured cells (whole population) and cells transfected with either CD44 (Figure 3A) or CD24 siRNAs (Figure 3B). CD44 and CD24 levels were significantly reduced in siRNA-transfected cells compared to the original population, thereby validating the specificity of our assay (Figure 3A-B). To identify miRNAs that play a role in enriching the CD44^high^/CD24^low^ cell population, we performed an unbiased functional screen to detect miRNAs capable of modulating the CD44/CD24 phenotype in HMECs (Figure 3C). Using a similar approach to our previous genome-wide siRNA screen for p16 modulators [35], we transfected actively proliferating cells (Passage 6, P6) with a library of 837 miRNAs, along with siRNA controls targeting siGLO (’cyclophilin B’; PPIB), CD44, or CD24. We established cut-off values to define miRNA hits based on the integrated intensity of CD44 and CD24 expression. The raw screening data and quantification for each phenotypic criterion are shown in Figure 3D. This strategy revealed that miR-106a-3p shifts primary cells toward a CD44^high^/CD24^low^ phenotype (Figure 3D). To validate these findings, we performed a secondary screen targeting the entire miRNA cluster (Figure 3E-F), retesting 28 miRNAs from the cluster using the same method as the primary screen (Figure 3C). Four miRNA hits scored a Z-score > 2 (Figure 3F), all of which induced a shift in the CD44^high^/CD24^low^ population (Appendix Figure S1A). The top hit was miR-106a-3p (Figure 3F). We then confirmed that miR-106a-3p induces the CD44^high^/CD24^low^ phenotype using flow cytometry based on the expression of CD44 and CD24 (Figure 3G). In cells transfected with the control mimic, only ∼10% exhibited the CD44^high^/CD24^low^ phenotype (Figure 3G). In contrast, cells transfected with the miR-106a-3p mimic demonstrated a fivefold increase in the CD44^high^/CD24^low^ phenotype, comprising approximately 50% of the total cell population (Figure 3G).

**Figure 3.**
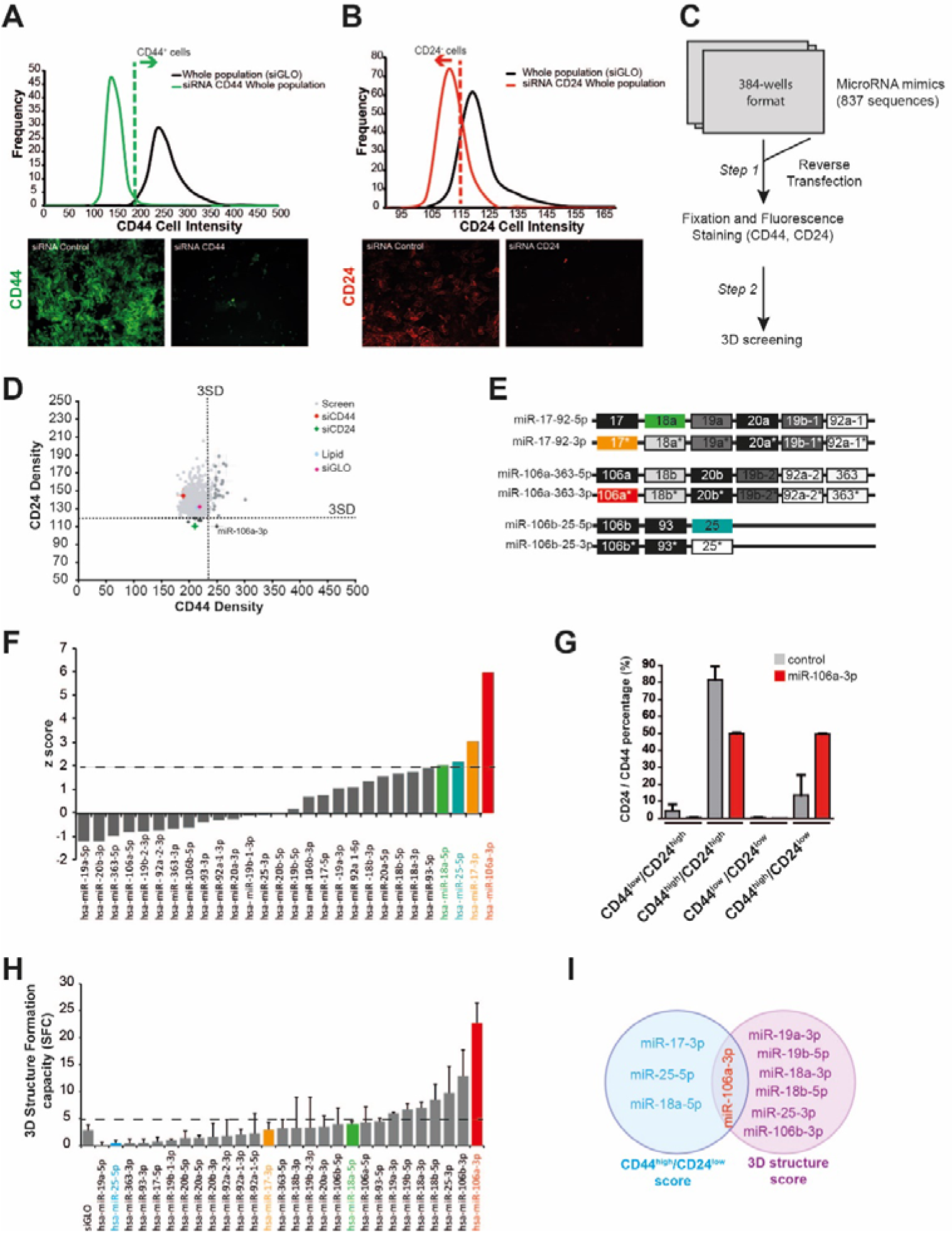
Identification of miR-106a-3p as the predominant miRNA in cells growing in 3D. **A,** Frequency distributions of CD44 intensity in HMECs at P6 (whole population, siGLO) as compared to HMEC-CD44-siRNA-depleted cells. Bottom panels are representative immunofluorescence pictures of HMECs stained with CD44 antibody (green): siGLO (left) and CD44 siRNA-knocked-down cells (right). **B,** Frequency distributions of CD24 intensity in HMECs at P6 (whole population, siGLO) as compared to HMEC CD24 siRNA knock down. Bottom panels are representative immunofluorescence pictures of HMECs stained with CD24 antibody (red): siGLO (left) and CD24 siRNA-knocked-down cells. **C,**Workflow for image-based miRNA screening for CD44^high^/CD24^low^ enhancers in primary human HMECs. HMECs were plated in 384-well plates and subjected to screening of miRNA libraries using optimized immunofluorescence staining for CD44 and CD24. **D,** Screening dot-plot showing the relationship between CD44 and CD24 intensities. Based on the frequency distributions generated for each phenotypic criterion (CD44 and CD24 intensity levels), we assigned highly stringent cutoffs for scoring positive hits in the genome-wide screen (dashed lines, 3 standard deviations (3SD) from the siGLO negative control. **E**, Members of the miR-17/92 cluster and its two paralogues miR-106a/363 and miR-106b/25. Red: miR-106a-3p; blue: miR-25-5p; green: miR-18-5p; orange: miR-17-3p. **F,** HMECs were transfected with miRNA mimics of the miR-17/92 cluster and its two paralogues and screened using conditions identical to the full screen. Z-Scores were calculated for individual miRNA mimics and plotted according to rank order. Dashed lines indicate 2 standard deviations (2SD) above the mean of the distribution. In colors are the miRNA above the 2SD. **G,** Mean percentages ± SEM of CD44/CD24 subpopulations from at least three independent sorting experiments in HMEC transfected with miR-106a-3p mimic as compared to control. **H,** 3D-SFC represents the mean numbers of 3D structures per cell seeded for each miRNA transfected in HMECs as compared to cells transfected with siRNA control (siGLO). Data are from three independent experiments. (mean±SEM). **I,** Venn diagram depicting the overlap of miRNA scoring in common between the CD44^high^/CD24^low^ and 3D structures scores. Note that the overall number of miRNAs in common would be the overlap of the intersect of these two Venn diagrams.

In parallel, to link these findings with the generation of stem/progenitor-like cells in 3D culture, we evaluated the frequency of 3D structure initiation after transfection with each of the 28 miRNAs from the miR-17/92 cluster (Figure 3E). Out of the 7 positive hits (Figure 3H), miR-106a-3p transfection showed the highest capacity for 3D structure formation (Figure 3H). Collectively, these results demonstrate that miR-106a-3p transfection induces two critical properties: 1) enrichment of CD44^high^/CD24^low^ cells and 2) enhanced initiation of 3D structures (Figure 3I).

### miR-106a-3p drives the generation of human 3D structures

To further investigate the role of miR-106a-3p, we generated retroviral vectors for miR-106a, as previously described [24], and evaluated its stable expression in 2D-cultured HMECs (Figure 4A-4B). First, we measured miR-106a-5p and miR-106a-3p expression using RT-qPCR in control cells (miR-Vector) and miR-106a-infected cells grown in 2D (Figure 4A). We observed that while miR-106a-5p was expressed in both infected groups, miR-106a- 3p expression was exclusive to miR-106a-infected cells, as confirmed by both RT-qPCR and in situ hybridization (Figure 4A-B). Additionally, when cells were cultured in 3D conditions, they expressed endogenous miR-106a-3p (Figure 4C). To confirm the physiological relevance of this miRNA, we used the DIANA-miTED database, which revealed that miR-106a-3p is expressed in various human tissues and cell lines (Appendix Figure S2).

**Figure 4.**
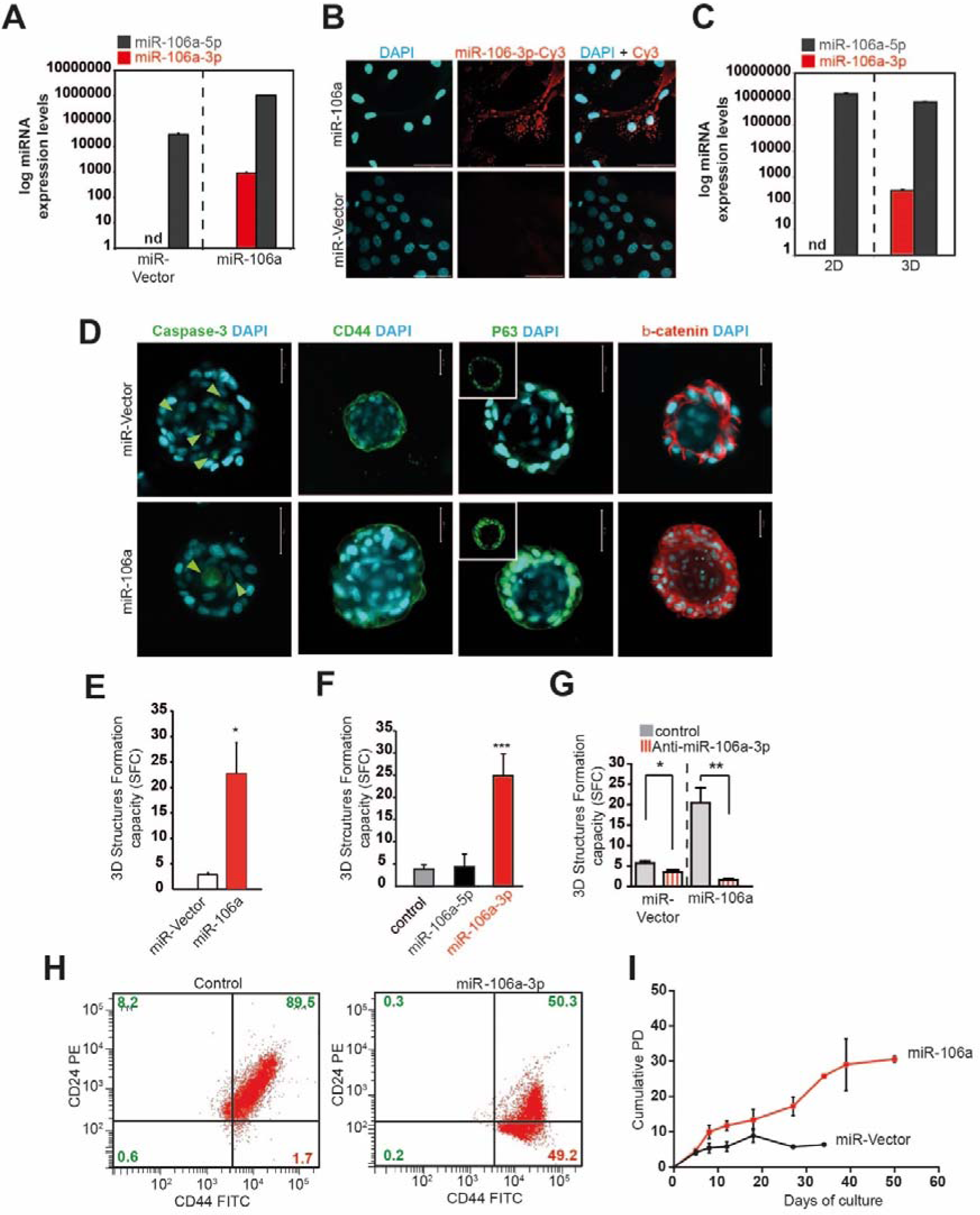
Properties of miR-106a-3p. **A,** Relative miR-106a-3p and miR-106a-5p expression levels determined by RT-qPCR in HMECs after retroviral infection with miR-Vector or miR-106a. n.d., not detectable. Data are from at least three independent experiments. (mean±SEM). **B,** FISH detection of miR-106a-3p in HMEC-miR106a stable cell lines. miR-106a-3p positive signals are visualized in red. Scale bar: 50 µm. **C,** Relative miR-106a-3p and miR-106a-5p expression levels determined by RT-qPCR in HMECs grown in 2D as compared to 3D; n.d., not detectable. Data are from at least three independent experiments. (mean±SEM). **D,** Confocal cross-sections of stable HMEC-miR106a 3D structures as compared to miR-Vector organoids stained with respectively active Caspase-3, CD44, p63 or β-catenin, and DAPI (blue) for nucleus. The arrows indicate apoptotic cells. Scale bars, 50 µm. **E** The bar graphs show the SFC, the mean number of 3D structures per well for miR-106a stable HMECs as compared to cells with miR-Vector. Statistical significance by Student’s t test is indicated by one (p < 0.05), two (p < 0.01), or three (p < 0.001) asterisks. Data are from at least three independent experiments. (mean±SEM). **F,** SFC represents the mean number of 3D structures per well in HMEC transfected with either control, miR-106a-5p or miR106a-3p mimics. Data are from at least three independent experiments. (mean±SEM). **G,** SFC as the percentage of 3D strcutures formed by cells seeded for either stable miR-Vector transfected with anti-miR control or stable miR-106a-HMEC transfected with anti-miR-106a-3p. Statistical significance by Student’s t test is indicated by one (p < 0.05), two (p < 0.01), or three (p < 0.001) asterisks. Data are from at least three independent experiments. (mean±SEM). **H**, Flow cytometric analyses of CD44/CD24 in HMEC transfected with either control or miR-106a-3p mimics. A minimum of 10,000 events were collected per sample. (n=3, mean ± SEM) **I,** Growth kinetics of human primary cells expressing either –miR-Vector or –miR-106a in long-term culture. The curve relationship between cumulative population doubling (PD) and duration of culture demonstrates a relatively linear PD rate with the progression of time. Data are from at least three independent experiments. (mean±SEM).

Next, we assessed the impact of miR-106a on 3D architecture using confocal microscopy. Immunofluorescence staining for Caspase-3, an apoptosis marker, demonstrated that miR-106a had no effect on luminal apoptosis (Figure 4D, Caspase-3). Furthermore, the well-defined cell/Matrigel interface and the myoepithelial layer, characteristic of 3D structures, remained intact despite miR-106a overexpression (Figure 4D, CD44 and p63). β- catenin, a marker of cadherin-based cell junctions, was also unaffected in terms of localization or cell junction integrity by miR-106a overexpression (Figure 4D, β-catenin). Taken together, these results indicate that miR-106a does not alter the morphogenesis of 3D structures and preserves their structural integrity and cellular interactions.

As expected, stable overexpression of miR-106a in primary HMECs markedly enhanced 3D structure formation capacity (Figure 4E). We then analysed the distinct functions of miR-106a-3p and miR-106a-5p in 3D structure-initiating cells by transfecting HMECs with either miR-106a-3p or miR-106a-5p mimics (Figure 4F). Overexpression of miR-106a-3p resulted in a fivefold increase in the number of 3D structures compared to both the control and miR-106a-5p (Figure 4F). In contrast, miR-106a-5p did not affect 3D structure formation, indicating the specific role of miR-106a-3p in maintaining this capacity (Figure 4F). Additionally, using LNA-anti-miR-106a-3p or LNA-control, we depleted endogenous miR-106a-3p levels and observed a 50% reduction in 3D structure-forming capacity, suggesting that miR-106a-3p is essential for 3D structure generation (Figure 4G). Flow cytometry analysis of miR-infected HMECs revealed that 89.5% of control cells (infected with miR-Vector) exhibited a CD44^high^/CD24^high^ phenotype, while only 1.7% displayed the CD44^high^/CD24^low^ phenotype (Figure 4H, left panel). In contrast, miR-106a-infected HMECs showed a significantly higher percentage (49.2%) of CD44^high^/CD24^low^ cells (Figure 4H, right panel). To evaluate whether this shift in population ratio affected cell survival, we examined the impact of miR-106a on population doubling in culture. Control cells (miR-Vector) stopped growing after approximately 10 population doublings, indicating limited proliferative potential. However, miR-106a extended the cells’ lifespan up to 30 doublings, suggesting a survival advantage (Figure 4I).

We then investigated whether miR-106a influences specific lineage sub-populations. Following lentiviral infection with either miR-106a-GFP or miR-Vector in primary HMECs, we analysed the heterogeneity of the miR-106a population via immunofluorescence. Cells expressing miR-106a-GFP (green) were mildly positive for cytokeratin 14 (CK14), a basal/myoepithelial marker, but negative for vimentin (Figure 5A). To determine if 3D structures generated in the presence of miR-106a exhibit enhanced self-renewal capacity compared to vector control, we evaluated serial passages of dissociated cells from single 3D structures. As shown in Figure 1D, most cells could generate a new 3D structure for at least one generation, and a few for up to three generations, indicating limited self-renewal potential. However, miR-106a expression enabled the same population to generate 3D structures for at least five generations, significantly extending their self-renewal capacity (Figure 5B).

**Figure 5.**
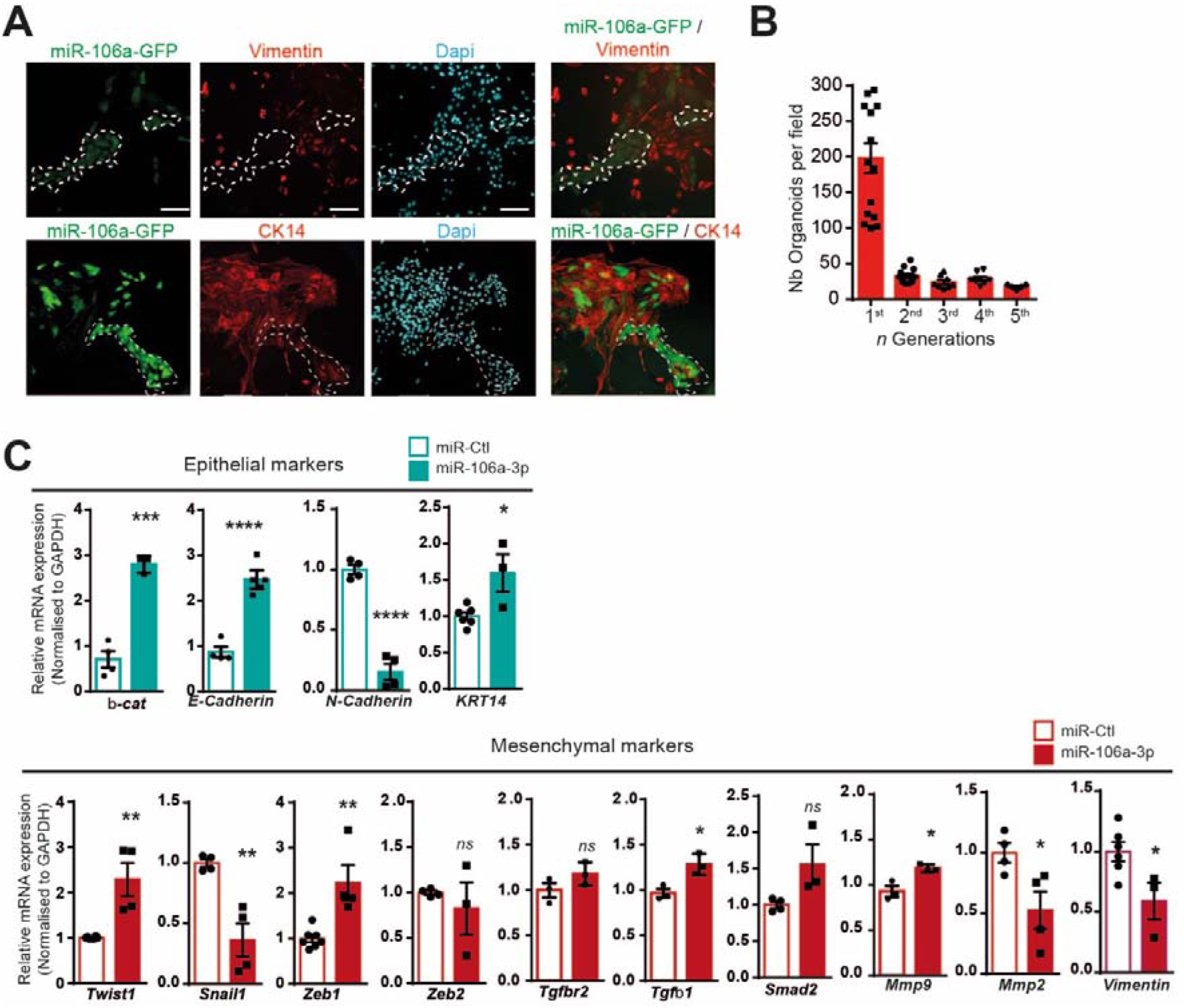
miR-106a-3p converts cell to a partial EMT-hybrid stade. **A,** Representative immunofluorescence (IF) analysis depicting loss of Vimentin expression stained for Vimentin (red), in miR-106a-GFP infected cells (green) as compared to non-infected cells, and nuclei with DAPI (blue). Additionally, basal/myoepithelial markers stained for CK14 (green), and DAPI. **B,** Quantification n generation of 3D structure development following miR-106a cell infection of cells. Data are presented as mean values ± SD. Data are from at least four independent experiments. (mean±SEM). **C**, Relative mRNA expression levels of key regulators of EMT determined by RT-qPCR in cells transfected with control miRNA (miR-Vector) or miR-106a-3p. In all graphs, means and standard errors are shown, and statistical significance by Student’s t test is indicated by one (p < 0.05), two (p < 0.01), or four (p < 0.0001) asterisks. Data are from at least three independent experiments. (mean±SEM).

Lastly, to investigate miR-106a-3p’s role in epithelial-mesenchymal transition (EMT), we performed RT-qPCR analysis on miR-106a-3p-expressing cells, as previously described,[2], targeting key EMT-related genes[36]. The epithelial marker set, including β-catenin, E-cadherin, N-cadherin, and KRT14, revealed significant upregulation of β-catenin, E-cadherin, and KRT14, while N-cadherin and vimentin were downregulated, confirming the observed vimentin-negative phenotype (Figure 5C, blue set). Among the mesenchymal markers tested—Twist1, Snail1, Zeb1, Zeb2, Tgfbr2, Tgfβ1, Smad2, MMP9, and MMP2—only Zeb1 and Twist1 were significantly upregulated, while Snail1 was downregulated (Figure 5C, red set). From the signalling group, Tgfβ1 showed a slight upregulation. Among non-genomic EMT processes, only MMP9 was upregulated.

In conclusion, these findings demonstrate that 1) the increased CD44^high^/CD24^low^ phenotype in miR-106a-3p-expressing cells drives the higher number of 3D structures; 2) miR-106a-3p induces a partial EMT phenotype; and 3) miR-106a-3p is essential for generating 3D structures, contributing to the maintenance of aSC/progenitor-like lineages.

### miR-106a-3p a key player of cell plasticity for differentiation

We performed transcriptomic analyses to determine whether miR-106a-3p influences cell plasticity, allowing the maintenance of adult stem-like cells or intermediate cellular states. To investigate the effect of miR-106a-3p on global gene expression, we extracted total RNA from miR-106a-3p-transfected HMECs and performed a microarray analysis using Affymetrix chips (HG-U133 Plus 2.0). Transfection with miR-106a-3p resulted in significant changes in the expression of 1,348 genes compared to the control mimic (Figure 6A). We analysed the transcriptomic data to investigate the differential activation of major signalling pathways and transcription factors. The major pathways are depicted in Figures 6B and 6C, revealing a significant downregulation of the MAPK, WNT, and PI3K pathways, while other oncogenic pathways, such as hypoxia response, JAK/STAT, and p53 pathways, were upregulated (Figures 6B and 6C). Additionally, we observed a reduction in the Wnt and PI3K signalling pathways, both known for their roles in maintaining stem/progenitor cells [37]. Moreover, SMAD4 and SMAD3 transcription factors were downregulated, correlating with the suppression of the TGFβ pathway. We also observed a differential regulation of genes involved in mammary breast development (Appendix Figure S4). To further understand the global transcriptional changes associated with miR-106a-3p, we compared our entire microarray dataset with established gene signatures through gene set enrichment analysis (GSEA). We identified an enrichment of downregulated gene sets involved in stem cell differentiation (Figure 6D). To explore the physiological role of miR-106a-3p in stem cell differentiation, we utilized human embryonic stem cells (hESCs), which express endogenous levels of miR-106a-3p (as observed in the DIANA-miTED database, Appendix Figure S2). Human ESCs differentiate more readily than aSCs. Throughout development, ESCs undergo epigenetic reprogramming, developmental patterning, and differentiation into various organs, each containing tissue-resident aSCs. Based on this, we hypothesized that miR-106a-3p in aSCs could be a remnant from ESCs. First, we evaluated the endogenous expression profile of miR-106-3p in hESCs (Figure 6E). Both miR-106a-5p, miR-106a-3p, and miR-302b (a marker of slowly-growing hESCs [38]) were detectable in hESCs (Figure 6E). Since hESCs are pluripotent and can indefinitely proliferate *in vitro* while maintaining the capacity to differentiate into all three germ layers (ectoderm, mesoderm, and endoderm) [39], we used hESCs to derive early stages of these germ layers (Figure 6F). We transfected hESCs with LNA-anti-miR-106a-3p (anti-miR106a-3p) or LNA-control (anti-miR-ctl) before initiating differentiation into endoderm (Figure 6G), mesoderm (Figure 6H), and ectoderm (Figure 6I). We then applied established protocols [40] to induce differentiation and determine whether blocking endogenous miR-106a-3p would alter transcriptional levels of OCT4, SOX2, and NANOG. In cells transfected with anti-miR-106a-3p, SOX2 expression decreased during differentiation into all three germ layers (Figure 6J, 6K, and 6L), while OCT4 expression decreased during endoderm and ectoderm differentiation (Figure 6H and 6J). NANOG expression, however, increased during mesoderm differentiation (Figure 6I). To further investigate the impact of miR-106a-3p depletion on hESC differentiation, we monitored the expression of specific genes upon inducing the three germ layers (Figure 6M, 6N, and 6O). Endodermal gene expression was minimally affected by miR-106a-3p levels (Figure 6M), while mesodermal and ectodermal gene expression increased upon miR-106a-3p downregulation (Figure 6N and 6O). These data demonstrate that miR-106a-3p plays a role in the early differentiation processes into the three germ layers, particularly the mesoderm and ectoderm layers.

**Figure 6.**
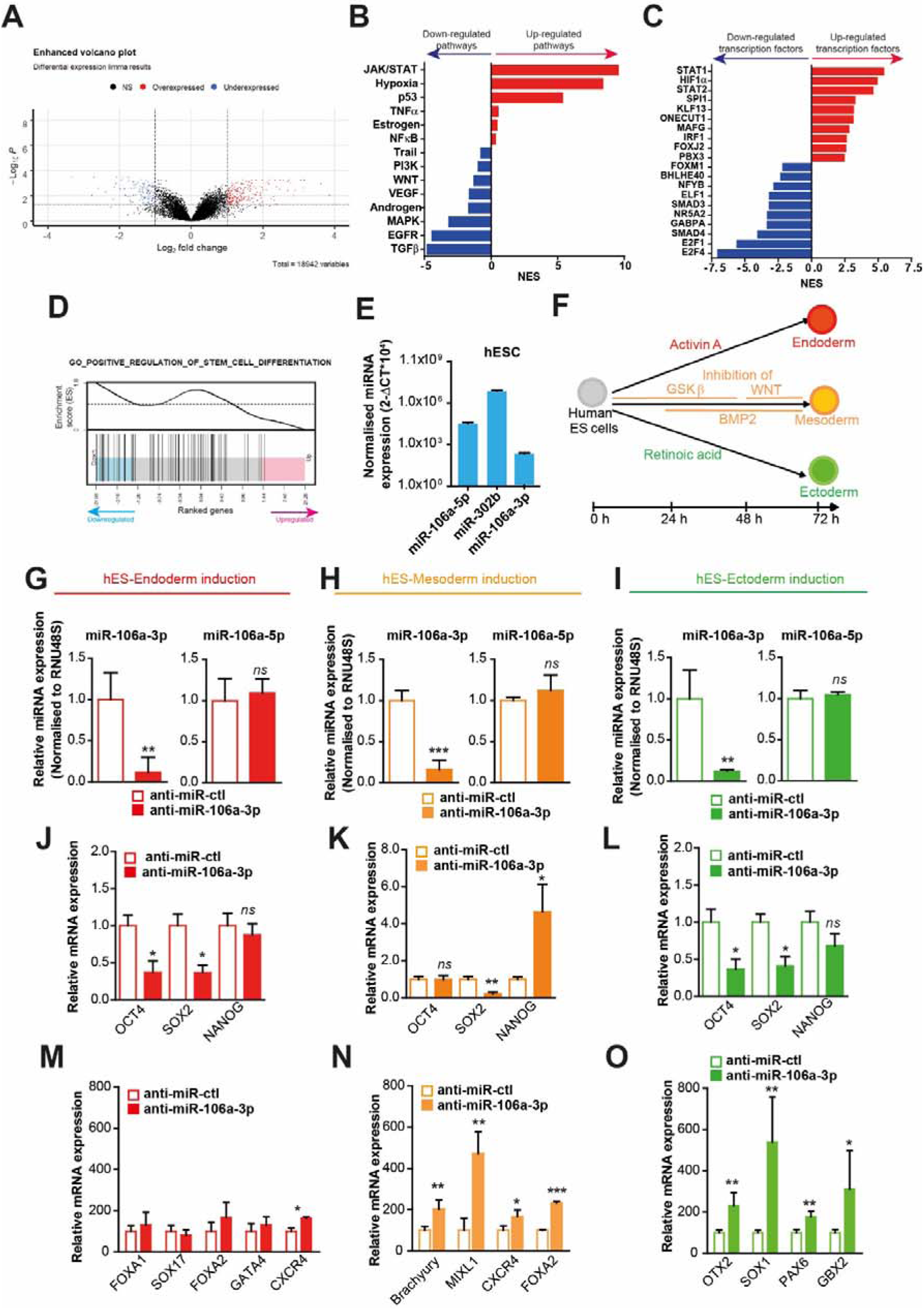
miR-106a-3p targets involved in downregulation of stem cell differentiation. **A,** Volcano plot comparing the gene expression fold changes and p-values between HMECs transfected with control mimic or miR-106a-3p, using the R package Enhanced Volcano. The plot shows the distribution of the total number of 18,942 genes in the core gene category were tested. Genes with fold change > 1 and FDR value < 0.05 are indicated in red, and genes with fold change < −1 and FDR value < 0.05 are indicated in blue. Data resulted in significant changes in the expression of 1,348 genes. **B-C**, Barplots illustrating 14 main pathways and the top-10 regulated TFs (up and down-regulated) following miR-106a-3p expression. The visualized numeric values are normalized enrichment scores (NES) of the top100 most responsive genes for each indicated pathway, whereas for the relative TF activities, the signed transcription factor (TF) - target gene interactions when utilized based on the DoRothEA database (weighted mean statistic). The relative molecular footprint alterations in control as compared to miR-106a-3p are represented by each bar, where NES < 0 represents down-regulation of specified pathway (B) or transcription factors (C). NES > 0 represents upregulation of specified pathway (B) or transcription factors (C). An absolute value of 2/-2 denotes the relative significance of the differential activity level. **D**, Barcode enrichment plot from the gene set enrichment analysis results, indicating down-regulation of stem cell differentiation pathway-related genes in the miR-106-3p-transfected group. The statistical significance of the enrichment is an FDR value of 0.024 and it is based on the multiple rotation gene-set testing (mroast) from the Limma R package (https://doi.org/10.1093/bioinformatics/btq401). **E,** Relative miR-106a-3p, miR-106a-5p and miR-302b expression levels determined by RT-qPCR in hESCs. Data are from at least three independent experiments. (mean±SEM). **F**, Schematic representation of the human ESCs differentiation process, including timeline and key signaling pathways that are modulated. **G-I**, Relative miR-106a-3p and miR-106a-5p expression levels determined by RT-qPCR in hESCs cells transfected with control mimic or anti-miR-106a-3p following induction of the endoderm (G), mesoderm (H) and ectoderm (I) differentiation. (n=3, mean ± SEM) **J-L**, Relative mRNA expression levels of key regulators of pluripotency in hESCs transfected with control mimic or anti-miR-106a-3p following induction of endoderm (J), mesoderm (K) and ectoderm (L) differentiation. In all graphs, means and standard errors are shown, and statistical significance by Student’s t test is indicated by one (p < 0.05), two (p < 0.01), or three (p < 0.001) asterisks. (n=3, mean ± SEM) **M-O,** Relative mRNA expression levels of differentiation target genes in hESCs transfected with control mimic or anti-miR-106a-3p following Endoderm induction (M), .Mesoderm induction (N) and Ectoderm induction (O). (n=3, mean ± SEM).

There is ongoing debate about the presence of unipotent, bipotent, or multipotent stem cells in mammary gland tissue [41–43]. The transcription factors OCT4, SOX2, and NANOG are known to maintain pluripotency in embryonic stem cells [44]. Over recent years, evidence has accumulated supporting the existence of stem cells in both mouse and human mammary tissue [45]. Various strategies, such as FACS sorting based on cell surface antigen expression, have been used to identify and isolate human breast stem/progenitor cells. Additionally, *in vitro* cell culture systems have been developed, allowing human mammary epithelial cells to proliferate in suspension as non-adherent mammospheres [30]. Simoe *et al.* demonstrated that stem cells isolated from both normal human breast and breast tumour cells express higher levels of embryonic stem cell genes, including NANOG, OCT4, and SOX2 [46]. Ectopic expression of these factors, particularly NANOG and SOX2, expands the stem cell population and enhances their ability to form mammospheres. Higher expression of NANOG, OCT4, and SOX2 was observed in CD44+CD24−/low and EMA+CALLA+ stem cell populations compared to other cell types. Overexpression of these genes led to an increase in stem cell populations, indicating their role in maintaining human mammary stem cells. Given the expression of *OCT4*, *SOX2*, and *NANOG* in hESCs and their presence in human breast cells, we hypothesized that aSCs derived from 3D structures might also express these genes. To test whether these transcription factors are involved in 3D structure initiation, we knocked down each gene and demonstrated their roles in 3D structure formation (Figure 7A). This suggests that *OCT4*, *SOX2*, and *NANOG* play significant roles in 3D growth. Furthermore, to determine whether these genes are induced during 3D differentiation, we monitored their mRNA levels over time in 3D cultures (Figure 7B). 3D structure development occurs in two phases: an initial phase where single cells proliferate to initiate 3D formation (around the first six days) and a proliferation arrest beginning around day 8 [47]. The expression of *OCT4*, *SOX2*, and *NANOG* peaked around day 6; subsequently, *OCT4* and *SOX2* levels decreased, while *NANOG* remained consistently high (Figure 7B). Finally, to investigate the potential modulation of these transcription factors by miR-106a-3p, we measured their mRNA (Figure 7C) and protein levels (Figures 7D-7F) in miR-106a-3p-expressing cells compared to controls. All three genes were upregulated in miR-106a-3p-overexpressing cells at both the transcriptional and post-transcriptional levels (Figure 7C-7F). This confirms that miR-106a-3p regulates *OCT4*, *SOX2*, and *NANOG* expression during 3D structure initiation.

**Figure 7.**
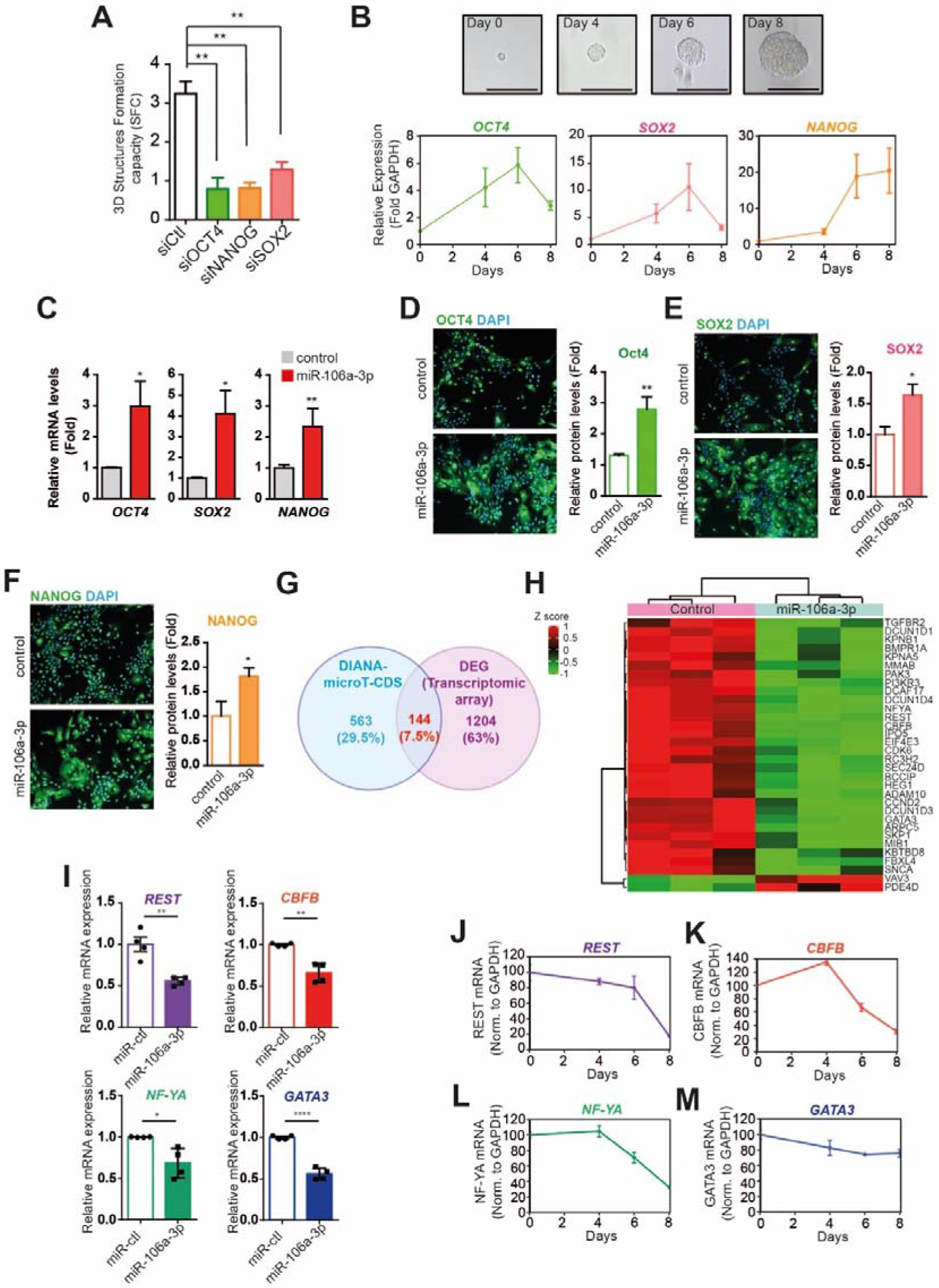
miR-106a-3p, a stem cell determining regulator for 3D structure development. **A,** SFC represents the percentage of 3D structure per seeded cell generated from HMECs transfected with siRNA control, siRNA-OCT4, siRNA-NANOG or siRNA-SOX2. (n=3, mean ± SEM) **B**, Relative *OCT4*, *SOX2* and *NANOG* expression levels determined by RT-qPCR in 3D growth of HMECs at Days 0, 4, 6, and 8. Data are from at least three independent experiments. (mean±SEM). **C**, Relative *OCT4*, *SOX2* and *NANOG* expression levels (measured by RT-qPCR and normalized to RNA48) following miR-106a-infected HMECs transfection with LNA-control (grey) or LNA-miR-106a-3p (red). In all graphs, means and standard errors are shown, and statistical significance by Student’s t test is indicated by one (p < 0.05), two (p < 0.01), or three (p < 0.001) asterisks. Data are from at least three independent experiments. (mean±SEM). **D-F**, Relative protein expression levels of OCT4 (D), SOX2 (E) and NANOG (F) determined by immunofluorescence analysis. Data are from at least three independent experiments. (mean±SEM). **G**, Venn diagram depicting the overlap of differentially expressed genes in the microarray experiment (1348 genes), with the predicted targets of the hsa-miR-106a-3p using the computational tool MicroT_CDS (http://diana.imis.athena-innovation.gr/DianaTools/index.php?r=MicroT_CDS/index) (707 genes), resulting in a number of 144 common genes. **H**, Heatmap of the 32 hub genes, resulted from the union of the REACTOME and GO biological processes, using the R package ComplexHeatmap **I**, *REST*, *CBFB*, *NFYA* and *GATA3* mRNA expression levels (measured by RT-qPCR and normalized to *RNA48*) following miR-106a-infected HMECs transfection with LNA-control (miR-ctl) or LNA-miR-106a-3p. Data are from at least three independent experiments. (mean±SEM). **J-M**, Relative *REST* (J), *CBFB* (K), *NFYA* (L) and *GATA3* (M) expression levels determined by RT-qPCR in 3D growth of HMECs at days 0, 4, 6, and 8. Data are from at least three independent experiments. (mean±SEM).

### Identification of miR-106a-3p targets through in silico analysis and transcriptomic profiling

In the last part of this study, we sought to identify the genes and mechanisms underlying the enhancement of 3D structure formation by miR-106a-3p. To determine if the significant expression changes observed after miR-106a-3p transfection (1,348 differentially expressed (DE) genes, as described in materials and methods) overlapped with data from predictive algorithms, we used the computational tool MicroT_CDS. This analysis identified 707 predicted targets of hsa-miR-106a-3p. The intersection of these datasets yielded 144 common genes (Figure 7G). Next, we conducted a functional enrichment analysis of the 144 common genes using REACTOME pathways and Gene Ontology, which led to the identification of 32 hub genes. An expression heatmap was generated to investigate the expression patterns of these genes within the microarray sample (see materials and methods). From this analysis, we identified 32 hub genes associated with miR-106a-3p, of which 30 were downregulated and 2 upregulated (Figure 7H). Among the downregulated genes, we identified 4 encoding transcription factors (*REST CBFB*, *NFYA*, and *GATA3*), which we subsequently examined in more detail (Figures 7I-7M). Transcript levels of these 4 putative targets were quantified by RT-qPCR in miR-106a-3p–transfected cells, revealing a statistically significant reduction in expression for each (Figure 7I). We then monitored the expression of these transcription factors during 3D structure development. *REST* (Figure 7J), *CBFB* (Figure 7K), and *NFYA* (Figure 7L) showed a significant decrease after 4 days of 3D structure development, while *GATA3* displayed more moderate changes (Figure 7M).

To validate the relevance of these 4 putative targets, we measured: 1) the impact on *OCT4*, *SOX2*, and *NANOG* mRNA expression (Figure 8B, 8E, 8H, and 8K), and 2) the frequency of 3D structure initiation (Figure 8C, 8F, 8I, and 8L). We hypothesised that siRNA-mediated knockdown of the four genes (*REST*, *CBFB*, *NFYA*, and *GATA3*) would impact *OCT4*, *SOX2*, and *NANOG* expression, thereby restoring the 3D structure-generating capacity observed after miR-106a-3p transfection. Each siRNA was first validated by its effect on the CD44/CD24 cell population (Appendix Figure S3), after which we assessed their impact on *OCT4*, *SOX2*, and *NANOG* expression. Knockdown of *REST* (Figure 8A) increased *OCT4* expression (Figure 8B), while knockdown of *NFYA* (Figure 8D) led to increased expression of both *OCT4* and *SOX2* (Figure 8E). *CBFB* knockdown (Figure 8G) resulted in increased *NANOG* expression and a moderate decrease in *SOX2* expression (Figure 8H), while *GATA3* knockdown led to higher *SOX2* and *NANOG* levels (Figure 8K). Finally, we demonstrated that knockdown of *REST*, *NFYA*, and *CBFB* could restore 3D structure formation, mimicking the effect of miR-106a-3p overexpression (Figure 8C, 8F, and 8I).

**Figure 8.**
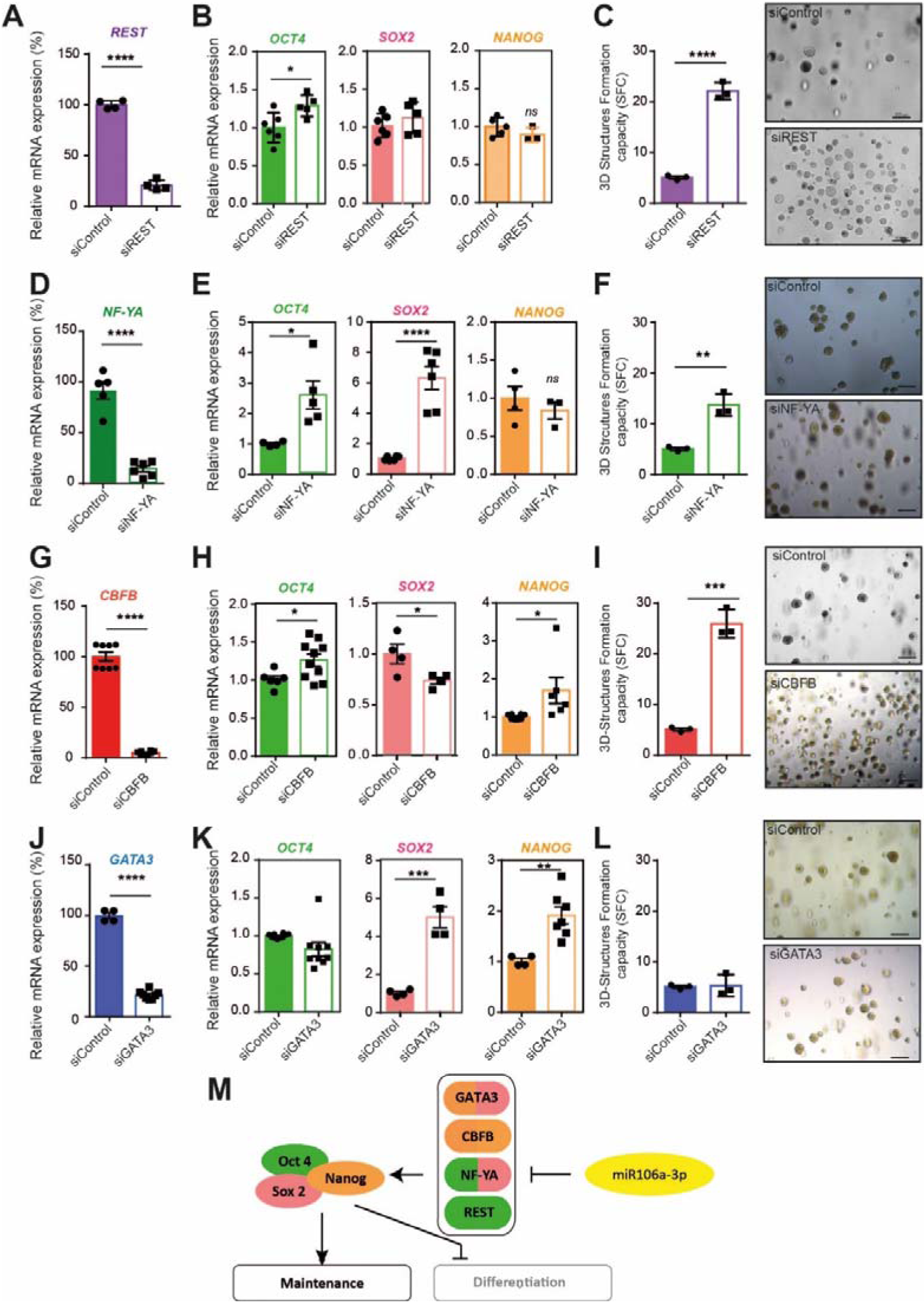
miR-106a-3p a barrier for cell differentiation program during mammary 3D structure initiation. **A,** Efficiency of *REST* gene silencing with specific human siRNA in transfected HMECs, as determined by RT-qPCR. Means and standard errors are shown. Data are from at least three independent experiments. (mean±SEM). **B**, Impact of *REST* silencing (siREST) on *OCT4*, *SOX2* and *NANOG* gene expression as determined by RT-qPCR. Data are from at least three independent experiments. (mean±SEM). **C**, SFC represents the percentage of 3D structures formed by seeded for control-transfected HMECs (siControl) as compared to siREST-transfected HMECs. Representative brightfield pictures of organoids from si-Control- and si-REST-transfected HMECs. Data are from at least three independent experiments. (mean±SEM). **D**, Efficiency of *NFYA* gene silencing with specific human siRNA in transfected HMECs as determined by RT-qPCR. Means and standard errors are shown. Data are from at least three independent experiments. (mean±SEM). **E**, Impact of *NFYA* silencing (siNFYA) on *OCT4*, *SOX2* and *NANOG* gene expression as determined by RT-qPCR. **F**, SFC represents the percentage of 3D structures formed by seeded for control-transfected HMECs (siControl) as compared to siNFYA-transfected HMECs. Representative brightfield pictures of 3D structures from control- and siNFYA-transfected HMECs. Data are from at least three independent experiments. (mean±SEM). **G**, Efficiency of *CBFB* gene silencing with specific human siRNA in transfected HMECs as determined by RT-qPCR. Data are from at least three independent experiments. (mean±SEM).**H**, Impact of *CBFB* silencing (siCBFB) on *OCT4*, *SOX2* and *NANOG* gene expression as determined by RT-qPCR. Data are from at least three independent experiments. (mean±SEM). **I**, SFC represents the percentage of 3D structures formed by seeded for control-transfected HMECs (si-control) as compared to siCBFB-transfected HMECs. Representative brightfield pictures of 3D structures formed from HMECs control- and siCBFB-transfected HMECs. Data are from at least three independent experiments. (mean±SEM). **J**, Efficiency of *GATA3* gene silencing with specific human siRNA in transfected HMECs as determined by RT-qPCR. Data are from at least three independent experiments. (mean±SEM). **K**, Impact of *GATA3* silencing (siGATA3) on *OCT4*, *SOX2* and *NANOG* gene expression as determined by RT-qPCR. Data are from at least three independent experiments. (mean±SEM). **L**, SFC represents the percentage of 3D structures formed by seeded for control-transfected HMECs (si-control) as compared to siGATA3-transfected HMECs. Representative brightfield pictures of 3D structures formed from HMECs control- and siGATA3-transfected HMECs. Data are from at least three independent experiments. (mean±SEM). **M,** Schematic representation of the mechanism initiated by miR-106a-3p. miR-106a-3p acts on 4 main transcription factors CBFB, NF-YA, GATA3 and REST. These transcription factors regulate the expression of OCT4, SOX2 and NANOG, which in turns favors stem cell maintenance and block cell differentiation.

## Discussion

While some argue that identifying stem/progenitor cells is unnecessary for primary tissue culture, understanding the stem cell niche is crucial for improving tissue-specific 3D structure growth conditions and sustaining 3D cultures indefinitely. In this study, we aimed to uncover the key factors promoting 3D structures derived from aSCs. This process involves identifying and characterising the cell populations initiating 3D structure formation, followed by an analysis of the transcriptional processes regulating 3D structure generation.

Here, we focused on miRNAs as key regulators capable of controlling both the generation and maintenance of 3D structures. Using an unbiased screening approach, we identified miR-106a-3p, a previously uncharacterised miRNA, as a master regulator of the stem/progenitor cell-like lineage that specifies the 3D structure-initiating cell population from human normal primary mammary epithelial cells. Our findings indicate that, in the experimental conditions explored, aSC-derived 3D structures are, at least in part, controlled by miR-106a-3p expression. Previous deep sequencing studies have shown that miRNA pairs (5p/3p) co-exist in roughly half of miRNA populations, often with varying concentrations of the 5p/3p species[48, 49]. Notably, the minor miRNA species—either 5p or 3p—exhibit evolutionary conservation in their seed sequences, underscoring their biological significance [50–52]. Several miRNAs have been found to either promote or inhibit stemness. For example, the loss of miR-205 expression induced a stemness phenotype in mammary epithelial cells, promoting EMT, altering cell polarity, and disrupting symmetric division by upregulating Zeb1/2 and Notch expression [53]. Additionally, miR-106a-3p has been linked to follicular lymphoma [54], gastric cancer [55], and renal carcinoma [56], highlighting its biological importance.

In breast tissue, EMT has been associated with stem/progenitor cell properties, including the expression of the CD44^high^/CD24^low^ profile. We hypothesised that miR-106a-3p may confer EMT features. However, instead of inducing a complete epithelial-to-mesenchymal transition, evidence increasingly suggests the presence of intermediate states where cells express both epithelial and mesenchymal traits [57]. Indeed, we observed that miR-106a-3p induced a hybrid EMT state, characterised by the expression of both epithelial markers (e.g., β-catenin and E-cadherin) and mesenchymal markers (e.g., TWIST1, ZEB1, TGFβ1, and MMP9). Moreover, cells expressing miR-106a-3p exhibited features associated with stemness, such as elevated TWIST1 levels. Interestingly, previous studies have shown that TWIST1 alone is not sufficient to induce EMT-related morphological changes [58, 59], suggesting that stem cell-like properties do not arise from the EMT programme itself but rather from a hybrid EMT state. ZEB1, another transcription factor upregulated by miR-106a- 3p, also plays a key role in the hybrid EMT state and the acquisition of stem cell-like characteristics [60]. Our findings suggest that stem/progenitor-like properties are linked to a hybrid EMT state rather than a complete epithelial or mesenchymal phenotype. Future studies should further investigate the time-frame window of miR-106a-3p induction during the EMT process.

Since major pathways such as Wnt, TGFβ/BMP, and Notch have been reported to be critical for maintaining mammary aSCs, thus we evaluated the potential involvement of these pathways. Stem cells possess self-renewal activities and multipotency, characteristics that tend to be maintained under hypoxic microenvironments;[61] therefore, it is not surprising to observe an up-regulation of hypoxia pathways and activation of HIF1α transcription factor. Remarkably, it has been shown by cell sorting that cells bearing a stem cell phenotype (CD44^high^/CD24^low^) express a constitutive activation of JAK-STAT pathway,[62] which we observed to be up-regulated along with STAT1 and STAT2 transcription factors. In parallel, we observe a down-regulation of Wnt signaling as well as PI3K pathways. Wnt is known to play an important role in stem cells maintenance; its inhibition has been shown to lead to the inactivation of PI3 kinase signaling pathways to ensure a balance control of stem cell renewal.[37] Moreover, in our model, we observed a down-regulation of SMAD4 and SMAD3 transcription factors, correlated with a down-regulation of the TGFβ pathway. Signals mediated by TGF-β family members have been implicated in maintaining and differentiating various types of somatic stem cells. [63] Moreover, we identified p53 pathway, in mammary stem cells, p53 has been shown to be critical to control the maintenance of a constant number of stem cells pool.[64, 65]

As cells differentiate and assume a specific lineage identity, some transcriptional mechanisms must switch on or off particular genes. To date, it remains unclear whether a defined transcriptional programme governs the aSCs maintenance. aSC identity, plasticity, and homeostasis are regulated by epigenetic and transcriptional networks specific to each lineage. By integrating gene arrays with miRNA/siRNA screening approaches, we identified 4 transcription factors (*CBFB*, *REST*, *NFYA*, and *GATA3*) as targets of miR-106a-3p, which are crucial for the generation of 3D structure-initiating cells and the maintenance of stem cell self-renewal (Figure 8M). Interestingly, knocking-down each of these genes phenocopied the effects of miR-106a-3p overexpression by modulating both the three core transcription factors (*OCT4*, *SOX2* and *NANOG*) and the 3D structure initiating cell population.

REST is a transcription factor expressed in epithelial cells [66], and plays a role OCT4/SOX2/NANOG transcriptional network [67]. It shares numerous target genes with OCT4, SOX2, and NANOG, several of which encode essential factors for cell maintenance in ESCs [67]. It binds a 21-bp DNA recognition sequence and has two repressor domains that recruit co-repressor complexes. REST binding sites in ESCs overlap with genomic regions that carry Polycomb-repressed chromatin in FACS-purified multipotent progenitors of the early embryonic pancreas [68]. Herein, REST also restrains differentiation into breast progenitors to favor aSCs maintenance in breast epithelial cells.

Core binding factor (CBF) is a heterodimeric transcription factor complex composed of a DNA-binding subunit, one of three runt-related transcription factor (RUNX) factors, and a non-DNA binding subunit, CBFB.[69, 70] There is only one CBFB subunit while the other subunit is encoded by three mammalian genes: *RUNX1*, *RUNX2*, and *RUNX3*, all of which require CBFB for their function. The targeted inactivation of CBFB abrogates the activity of all RUNX complexes[71]. Targeted knock-out of RUNX genes has revealed distinct roles for these proteins in development, with RUNX1 being required for hematopoiesis[72], RUNX2 for osteogenesis,[73] and RUNX3 for neurogenesis and the control of gastric epithelial cell proliferation.[74, 75]. The RUNX/CBFB complexes have been shown to play a role in the stem cells maintenance by activating FGF signaling loops between the epithelium and the mesenchyme [71]. Additionally, CBFB has been found to be upregulated in the lactating mammary gland and is necessary for lobulo-alveolar development [76]. Furthermore, CBFB knockout has been shown to lead to a down-regulation of SNAIL and VIM expression during the hybrid-EMT, indicating a more epithelial state in breast cell lines [77]. These data confirm our observations, indicating that miR-106a-3p, through the down-regulation of CBFB, induces a down-regulation of both SNAIL1 and VIM in our model. Therefore, it is not surprising that controlling CBFB through miR-106a-3p expression participates in the maintenance of aSCs.

NF-Y, a ubiquitously expressed trimeric transcription factor, has a dual role as an activator and repressor of transcription [78]. The heterodimer protein complex comprises three subunits (NF-YA, NF-YB, and NF-YC). NF-YA is considered as the limiting regulatory subunit of the trimer, since it is required for complex assembly and sequence-specific DNA binding. NF-Y has previously been identified as a marker of CSCs in hepatocellular carcinoma and embryonic carcinoma cells [79–81]. In addition, it has been shown to regulate the expression of several human *SOX* genes, including *SOX2* [82], NF-Y also regulates the expression of several human SOX genes, including *SOX2*[82], *SOX9* [83], and *SOX18*[84].

This transcriptional activation function of NF-Y is mediated, at least in part, by direct binding to CCAAT boxes within promoters of target genes and by making complex interplay with other factors involved in transcriptional regulation of human *SOX* genes. It has also been shown that the NF-Y binding site CCAAT within the proximal region of the human *SOX2* gene promoter plays a key role in regulating SOX2 expression in cervical CSCs, establishing that NF-YA is essential for maintaining CSCs characteristics. Interestingly, NF-Y has been shown to regulate ATF6 expression [85] which is involved in protein homeostasis. Recently, it has been proposed that cell proteostasis restrains protein synthesis for the maintenance of stem cells [86]. As a consequence, we can speculate that NF-YA might also participate in stem cell maintenance through the regulation of cell proteostasis. Herein, we found that NF- YA regulates the expression of SOX2 for the maintenance of breast aSCs, consistent with the literature.

The GATA transcription factors play critical roles in the gene regulatory networks governing cell fates specification and maintenance. Among them, the zinc finger transcription factor GATA-3 is known to serve as a key regulator of commitment and maturation of the luminal breast epithelial lineage [87] and displays essential roles in the morphogenesis of the mammary gland during embryonic and adult stages. Notably, GATA-3 has been established to be a critical regulator of luminal differentiation, and its deficiency leads to an expansion of luminal progenitors and a concurrent block in differentiation [87]. These data confirmed our observations that miR-106a-3p could act on differentiation through the down-regulation of the GATA3 transcription factor.

It is now well recognized that the lineage hierarchy of the mammary epithelium is composed of a basal/myoepithelial lineage that can function as multipotent mammary stem cells, capable of generating multilineage functional mammary epithelia *in vivo* [43, 88]. These cells have thus been viewed as the primary hierarchy of the mammary epithelium, giving rise to more restricted lineage-specific progenitors, and being responsible for the continuous generation of all mammary epithelial lineages in adults. However, it remains unclear whether cells possessing mammary stem cell potential in the basal layer are distinct from differentiated, non-stem cell basal cells or if stem cell potential is a general feature of cells present in this layer.

The luminal compartment has been shown to contain mature differentiated luminal cells, unable to generate 3D structures, as well as a subpopulation of luminal progenitor cells capable of giving rise to structures containing luminal cells [43, 88]. We thus performed lineage-tracing experiments using tissue dissociation and surface marker staining and sorting to identify three main populations: mature luminal cells, luminal progenitors, and basal/stem cells. We found that cells growing in 3D are enriched in luminal progenitors and miR-106a enriches this same population. Interestingly, it has been shown that these luminal cells are also able to give rise to ductal structures containing both lineages [89], indicating that a more limited mammary stem cell potential may be present in this lineage as well. It has been suggested that mammary stem cells might possess lineage characteristics that are at an intermediate point on the basal–luminal axis. Indeed, fetal mammary stem cells (fMaSCs), which give rise to the entire mammary gland, do indeed appear in fact to express both luminal and basal markers, representing an intermediate position on this axis, and their gene expression signature is closer to that of adult luminal progenitors than adult basal cells [41, 90].

Mechanistically, miR-106a-3p targets a specific set of genes, namely *CBFB, REST*, *NFYA* and *GATA3*, to finely regulate the expression of OCT4, SOX2, and NANOG, thereby reducing heterogeneity within the 3D structure-initiating cell population (Figure 8M). Consequently, a complex mechanism is clearly established to finely modulate the expression of key transcription factors in 3D culture, a process conserved throughout development in adult and embryonic stem cells. Recent reports have demonstrated that the differentiation of human aSCs [91] and mouse ESCs [92] is modulated through post-transcriptional attenuation of key factors such as OCT4, SOX2, and NANOG . A recent study by Doffou *et al.*, has also reported a similar phenomenon in an organoid hepatocyte model, showing that OCT4 expression is induced, starting at day 6 [93]. This induction is seen in hepatocytes in anticipation of trans-differentiation. This increase of OCT4 could be attributed to differentiation in our model, based on previous studies showing that both under- and over- expression of OCT4 could lead to cell differentiation [94].

miRNA-directed regulations provide a way to finely tune aSCs’ self-renewal and differentiation. Indeed, miRNAs play an important role in gene regulation for ESCs’ pluripotency, self-renewal, and differentiation. These miRNAs can be divided into two subgroups: pluripotency-inducing miRNAs and pro-differentiation miRNAs. The first subgroup, including miR-137, miR-184, miR-200, miR-290, miR-302, and miR-9, is exclusively expressed during the pluripotent state and rapidly decreases upon differentiation stimuli [17, 18]. By contrast, pro-differentiation miRNAs, such as let-7, miR-296, miR-134, and miR-470, regulate the differentiation processes in pluripotent cells [38].These miRNAs are upregulated during ESCs differentiation and inhibit the expression of pluripotency factors, including NANOG and SOX2 [38]. Even though aSCs are epigenetically and temporally very distant from ESCs, we found that the miR-106a-3p impaired differentiation in ESCs and 3D- structure formation in aSCs suggesting that this miRNA might be a remnant of ESCs. Our data show that miR-106a-3p features as a miRNA participating in the modulation of the core factor network (OCT4, SOX2, and NANOG), which in turn inhibits differentiation and favors maintenance of stem/progenitor cells (Figure 8M). Indeed, miR-106a-3p is sufficient by itself in targeting a specific set of genes *CBFB, REST, NF-YA,* and *GATA3*, and to induce expression of *OCT4*, *SOX2*, and *NANOG*. Finally, the role of miR-106a-3p is of particular interest in explaining how mammary epithelial cells acquire stem cell-like properties in normal conditions. Indeed, the capacity of miR-106a-3p to promote stem cell-like behavior in 3D gives us some clues on how the stem cell status may be specified in mammary cells. Our study underscores the cooperative action of transcription factor networks in establishing stem/progenitor cell identity, providing comprehensive regulatory principles for human epithelial homeostasis. To date, it was not known whether a transcriptional program existed to maintain aSCs, a question addressed in this study (Figure 8M). Understanding the transcriptional networks and signalling pathways involved in 3D structure formation could enhance the use of organoids as models for studying organ function, disease, and therapy.

## Acknowledgments

We gratefully acknowledge the members of the ARTiSt Lab for their critical remarks. FD was supported by grants from the “Ligue Contre le Cancer Gironde” and from the “Site de recherche intégrée sur le cancer de Bordeaux” (SIRIC Brio). This work has been supported by grants from the “Région Nouvelle-Aquitaine” (DF).

## Author Contributions

Conceptualization, D.F., F.D., G.L., and A.C; Methodology, D.F., F.D., and M.P.; Investigation, D.F., F.D., E.V, and M.P.; Writing – Original Draft, D.F.; Writing – Review & Editing, D.F., F.D., J.R., E.V and M.P; Resources, D.F. and M.P.; Supervision, D.F., and M.P.

## Materials and Methods

### Contact for reagent and resource sharing

Further information and requests for resources and reagents may be directed to and will be fulfilled by the Lead Contact, Delphine Fessart.

### Experimental model details

#### Cell lines

Normal HMECs with finite life-span have been previously fully described [24, 35]. Finite-lifespan HMECs were provided by the Human Mammary Epithelial Cell (HMEC) Bank from M. Stampfer. M87A-type media was used [19]. M87A media has been reported to support pre-stasis HMEC growth for as much as 60 population doublings (PD), and to maintain luminal cells for as many as 8 passages (∼30 PD). It has been shown that M87A supports pre-stasis HMEC growth, including myoepithelial, luminal, and progenitor cells for 30–60 PD [95]. HUES cells (HUES9) were cultured as previously described [40].

#### Cell culture

For standard 2D culture, primary human mammary epithelial cells at passage 4 were established and maintained in M87A medium as previously described [24]. HMECs cells at Passage 6 (P6) were used for the miRNA screening and follow-up miRNA studies, unless otherwise stated. For three-dimensional culture (3D), cells were grown in laminin-rich basement membrane growth factor-reduced Matrigel (BDBiosciences) (Matrigel) as we previously described.^5^ Mycoplasma testing was performed prior to all experiments in this study.

## Methods

### High-content miRNA screening

The miRNA screen was performed in triplicate, using the Human pre-miR miRNA library (Ambion), consisting of 837 miRNAs, together with control small interfering RNAs (siRNAs) targeting Cyclophilin B (Dharmacon), CD44, and CD24 (Qiagen). HMECs at P6 were reverse-transfected with 30 nM miRNA in 384-well format using HiperFect (QIAGEN), in triplicate using Janus apparatus (Perkin Elmer) of the POETIC plateform. Plates were incubated for 48 h, medium was changed and fixed/stained 72 h later with CD44-FITC conjugated antibody (Abcam), CD24 antibody (BD Biosciences) and GtαMo AlexaFluor546 (Invitrogen), 4′,6-diamidino-2-phenylindole (DAPI, Sigma). For the screening in 3D culture, cells were fixed following 8 days after transfection. High-content images were acquired with the Cytation3 (Bioteck) at 4× magnification, and analysis was performed using the Analysis software (Bioteck). The Z-score provides a metric of the median absolute deviation by which an individual miRNA transfected condition (averaged over three replicates) differs from the population median (median percentage CD44^high^/CD24^low^ population).

### Flow cytometry

Following trypsinization, cells were strained through a 40 μm nylon mesh to ensure single cells are obtained and suspended in ice-cold solution to obtain a density of 1 × 10^6^ cells/ml. Antibodies (CD44 conjugated with FITC (Abcam, Ab19622); CD24 conjugated with phycoerythrin, PE (BD Biosciences, 555428); CD-24 (BD Biosciences, 55046); CD49f conjugated to Cy5 (BD Biosciences, 551129) were added to the cell suspension at concentrations suggested by the manufacturer and cells were incubated at 4°C in the dark for 45 min. These labeled cells were washed twice, suspended in PBS and analysed using a flow cytometer (Becton Dickinson). The cells were stained with either isotype-matched control antibodies or with no primary antibody as negative controls. No difference was observed between these two controls. To separate the ALDH positive population by FACS, the ALDEFLUOR kit (StemCell Technologies) was used. Cells were suspended in ALDEFLUOR assay buffer containing ALDH substrate (BAAA, 1 μM per 1×10^6^ cells) and incubated during 40 minutes at 37°C. As negative control, for each sample of cells an aliquot was treated with 50mM diethylaminobenzaldehyde (DEAB), a specific ALDH inhibitor.

### RNA isolation and miRNA microarray

Total RNAs were isolated from three independent samples of HMEC-transfected cells using the miRNeasy Kit (Qiagen) according to the manufacturer’s instructions. The quantity and size of RNAs were analysed for concentration, purity and integrity by using spectrophotometric methods in combination with the Agilent Bioanalyzer (Agilent Technologies).

Microarray analyses were performed on 3 independent replicates of mimic control transfected cell samples (control), 3 independent replicates of miR-106a-3p transfected cell samples.

Complete gene expression analysis was performed with R (R version 3.6.1)/Bioconductor software (https://doi.org/10.1038/nmeth.3252). Initially, the raw data were imported with oligo R package (1.48.0) (https://doi.org/doi:10.18129/B9.bioc.oligo) and processed (background subtraction, quantile normalization and summarization with median polish method) using the RMA algorithm (https://doi.org/10.1093/biostatistics/4.2.249). In addition, for the annotation process, the R packages affycoretools (1.56.0) and hgu133plus2.db (3.2.3) were used to map probe sets to gene symbols (https://doi.org/doi:10.18129/B9.bioc.affycoretools). Next, after the removal of control features, a non-specific intensity filtering procedure was applied to remove probesets that were not expressed at least in one of the two conditions (control or transfected samples). Finally, aiming to identify differentially expressed genes between transfected and non-transfected samples, linear models were fitted and statistical inference was estimated using the limma R package (3.40.6) (https://doi.org/10.1093/nar/gkv007). Differentially expressed genes were identified using an FDR value < 0.01 & an absolute value of log2- fold change > log2(1.5). Regarding the visualization part of the differential expression analysis, volcano plots were created with the EnhancedVolcano R package (1.2.0) (https://doi.org/doi:10.18129/B9.bioc.EnhancedVolcano), whereas the R package ComplexHeatmap (2.0.0) was utilized for the creation of the gene expression heatmaps based on selected gene signatures (https://doi.org/10.1093/bioinformatics/btw313). The gene expression data have been deposited in the ArrayExpress database at EMBL-EBI (www.ebi.ac.uk/arrayexpress) under accession number E-MTAB-6594. The R code implemented for the analysis is available upon request. We have analysed our transcriptomic data, using the PROGENy R package (version 1.16.0) (https://doi.org/10.1038/s41467-017-02391-6) and DoRothEA (version 1.6.0)-decoupleR (version 2.1.6) computational pipeline (https://doi.org/10.1101/gr.240663.118, https://doi.org/10.1093/bioadv/vbac016), to investigate the differential activation of major signaling pathways and transcriptional factors.

### Functional enrichment analysis

To exploit the biological mechanisms involved in the miR-106a-3p transfection effect, the BioInfoMiner interpretation web platform was used (10.4018/IJMSTR.2016040103; https://doi.org/10.15252/emmm.201707929). BioInfoMiner implements an automated and robust network analysis of functional terms, by the integration of semantic information through different biomedical vocabularies, aiming to elucidate the significantly perturbed biological processes, and critical genes with centrality role affected in the studied phenotype (https://doi.org/10.1038/s41467-022-30159-0). Furthermore, in order to unravel if specific mechanisms related to stemness and differentiation are significantly altered in the transfected samples, a customized enrichment analysis approach was applied through rotation gene set tests (https://doi.org/10.1093/bioinformatics/btq401), using the limma R package (mroast function). For the gene set signatures, we initially selected from the Molecular Signatures Database (https://www.gsea-msigdb.org/gsea/msigdb/), (https://doi.org/10.1073/pnas.0506580102) the ontology and curated gene sets (version 7.1). Then, as a final step we kept only the terms/pathways that included the phrases “STEM” or “NOTCH” and had at least 10 genes in the respective signature. Enrichment analysis was performed using REACTOME pathways (https://doi.org/10.1093/nar/gkab1028) and Gene Ontology (https://doi.org/10.1093/nar/gkaa1113) to identify 32 hub genes, and generate an expression heatmap to investigate the expression patterns in the microarray sample.

### miRNA target identification

The miRNA targets predictions based on miRanda, DianaMT, miRDB and miRWalk were downloaded from www.microrna.org (August 2010 release), http://zmf.umm.uni-heidelberg.de/apps/zmf/mirwalk2/ and from http://mirdb.org/miRDB/. We used the computational tool MicroT_CDS (https://doi.org/10.1093/nar/gkt393)

### miRNA target stem cells signature analysis

Gene set stem cells enrichment analysis for predicted miRNA targets was carried out using the web interface of Stem checker (http://stemchecker.sysbiolab.eu/) using default settings. *miRNA and antigomiR transfections* HMECs were transfected with 30 nM miRNA or 30 nM antigomiR (anti-miRNA) in 384-well plates using HiperFect (Qiagen), and the protocol described above for ‘High-content miRNA Screening’ was followed. For siRNA transfections, pools of three siRNA per target were purchased (Qiagen) and the cell were transfected with Hyperfect (Qiagen) according to manufacturer instructions.

### Quantitative reverse transcriptase-polymerase chain reaction

Methodology for quantitative reverse transcriptase-polymerase chain reaction (RT-qPCR) has been described previously. Quantitative RT-PCR reactions were performed with SYBR Green Master Mix (ABI). For siRNA knockdown experiments, three siRNA per targets were used and, RNA was extracted from 1× 10^5^ cells 48hr post-transfection. *GAPDH* levels were quantified for each cDNA sample in separate qPCR reactions and were used as an endogenous control. Target gene-expression levels were quantified using target specific probes. Values were normalized to the internal *GAPDH* control and expressed relative to *siGLO* transfected control levels (100%). All qPCR reactions were run in triplicate from three independent samples. Primers used for qPCR are in Appendix Table S1.

### Retroviral stable cell lines

106a-5p/-3p miRNA hit was cloned into MirVec as previously described.^3^ After sequence verification, 5 mg of plasmid DNA was transfected into HMEC P5 was transduced into Phoenix packaging cells using Fugene (Roche, Basel, Switzerland). Viral supernatant was harvested 48 h after transfection. Target HMECs were seeded in a six-well plate at a density of 5000 cells/cm^2^ and spinfected the following day at 32 °C, 350 r.p.m. for 1 h with viral supernatant in the presence of 8 mg/ml polybrene. Cells were selected with blasticidin (3 mg/ml). Cells were harvested for RT-qPCR analysis.

### Immunofluorescence

Fixed cells were permeabilized with 0.1% Triton X-100 (Sigma) for 30 min at room temperature (RT) cells were stained for 2 h at RT with a primary antibody followed by a secondary antibody staining for 1 h at RT (AlexaFlour-488-conjugated goat anti-mouse antibody (Invitrogen). Cells were imaged on Leica Dmi8 microscope. Images were analysed using Leica software. Primary antibodies used were CD44 (BD Biosciences, 550392); CD44 (Abcam, Ab19622), CD24 (BD Biosciences, 550426), cleaved Caspase-3 ((Asp175), Cell Signaling, 9664S), beta-catenin (BD Biosciences, 610153), p63 (clone 4A4; Santa CruzBiotechnology, sc8431), MoαCK14 (Ozyme), CK18 (Santa CruzBiotechnology, sc32722), EpCAM (EBA-1; BD Biosciences, 347197); MUC1 (Ozyme, BTMBNC80960); CK5 (Abcam, Ab17130), Vimentin (Sigma, V6389). Secondary antibodies were the appropriate AlexaFluor-488 or AlexaFluor-546 antibody (Invitrogen). DAPI and CellMask Deep Red (Invitrogen) were also included. Images were collected with the Dmi8 microscope (Leica) or the Zeiss 510 Meta Confocal microscope (Zeiss) and Developer Software (Leica) used for image analysis.

### In situ hybridization (ISH) and microscopy

ISH was performed by using specific DIG-labeled miRNA LNAprobes from Exiqon. Briefly, cells were fixed in 4% paraformaldehyde for 30 min, followed by 70% ethanol for at least 16 h at 4°C. Cells were then permeabilized with 0.1% Triton X-100 for 10 min. The washed cells were then pre-hybridized with a prehybridization buffer (46 SSC, 25% formamide, Denhardt’s solution, 2% blocking reagents, 0.25 mg/ml yeast tRNA, 0.25 mg/ml salmon sperm DNA) for 30 min at room temperature, followed by hybridization at 23 °C below the Tm of the LNA probe for 2 h. The cells were subsequently washed with Washing Buffer I (2X SSC with 0.1% Tween 20), II (2X SSC), and III (0.5X SSC) at the hybridization temperature. Cells were blocked with a signal enhancer (Lifetechnologies) for 1 h at room temperature, and then incubated with a mouse anti-DIG antibody at a dilution of 1:1000 at 4°C overnight. Cells were washed with PBS three times to remove unbounded mouse anti-DIG antibody. Then, cells were incubated with a fluorescently labeled secondary antibody. To confirm that the ISH signals were indeed from the specific hybridization of the probes with the target RNA, the cells stained with a specific miR-scramble DIG-labeled miRNA LNAprobes from Exiqon. The DNA was stained with DAPI. The samples were mounted on a fluorescent mounting medium (Dako). The images were taken with a LSM-510 Meta (Zeiss) confocal microscope.

### ESCs differentiation

HUES cells (HUES9) were cultured as previously described [40]. Endoderm was induced by treating the cells for 3 days with 100 ng activin A (Peprotech, France) in DMEM supplemented with 10% FCS. Mesoderm was induced by culturing the cells in RPMI supplemented with 20% B27 (Thermofisher, France) and added with 5 µM CHIR 99021 (Stem cell, France) for 24 hr, then with BMP2 (10 ng/ml, Thermofisher, France) and CHIR 99021 (5 µM) the second day and finally IWR1 (2 µM) and BMP2 (10 ng/ml) the third day. Ectoderm was induced in RPMI supplemented with N2 medium (Thermofisher) and 0.5 µM retinoic acid for three days.

### Quantification and statistical analyses

Quantification data are presented as means ± SEM. Statistical significance was analysed using an unpaired Student’s t test. A difference at p < 0.05 was considered statistically significant.

