## Appendix for "A miRNA screen identifies a transcriptional program controlling the fate of adult stem cell"

**Table Contents**

**Figure S1**: Scatter plots of CD44/CD24 intensity in normal HMEC transfected with miR-106a-3p ; miR-17-3p; miR-25-5p; and miR-18-5p.

**Figure S2.** miR-106a-3p expression in tissue and cell lines from DNA-miTED database.

**Figure S3.** CD44 and CD24 profiles following siRNA against REST, NF-YA, CBFB and GATA3

**Figure S4.** Genes identified as involved in Mammary Gland development

**Table S1** List of the primer sequences used in the study.

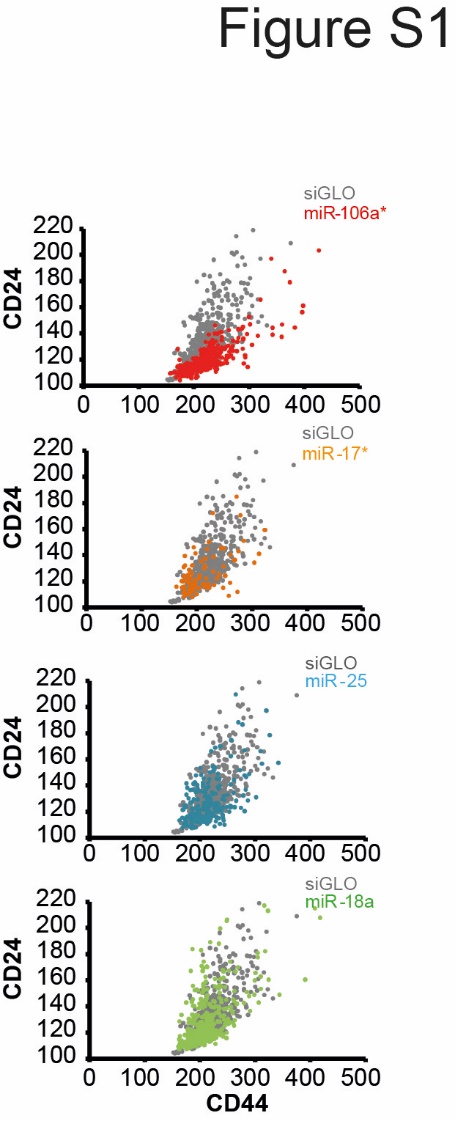

**Figure S1**

Scatter plots of CD44/CD24 intensity in normal HMEC transfected with miR-106a-3p (red); miR-17-3p (orange); miR-25-5p (blue); and miR-18-5p (green).

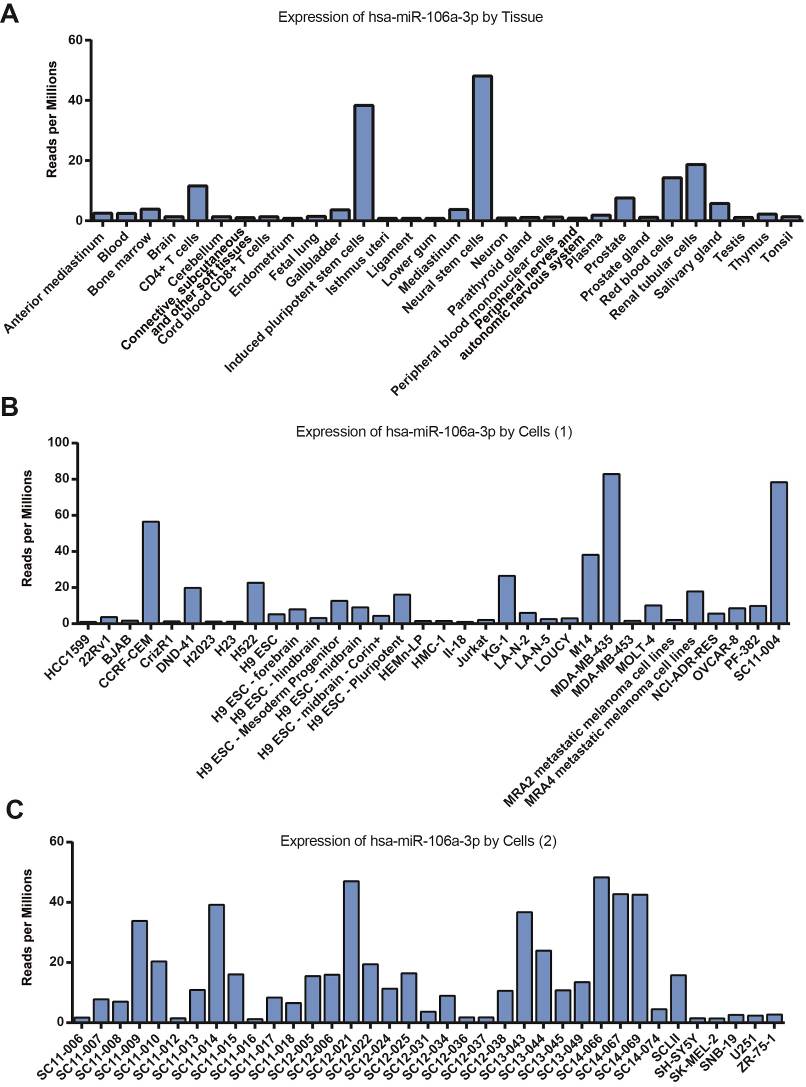

**Figure S2. miR-106a–3p expression in tissue and cell lines from the DIANA -miTED database.**

**A,** Mean expression of miR-106a-3p in various tissue. **B-C,** Mean expression of miR-106a-3p in various cell lines.

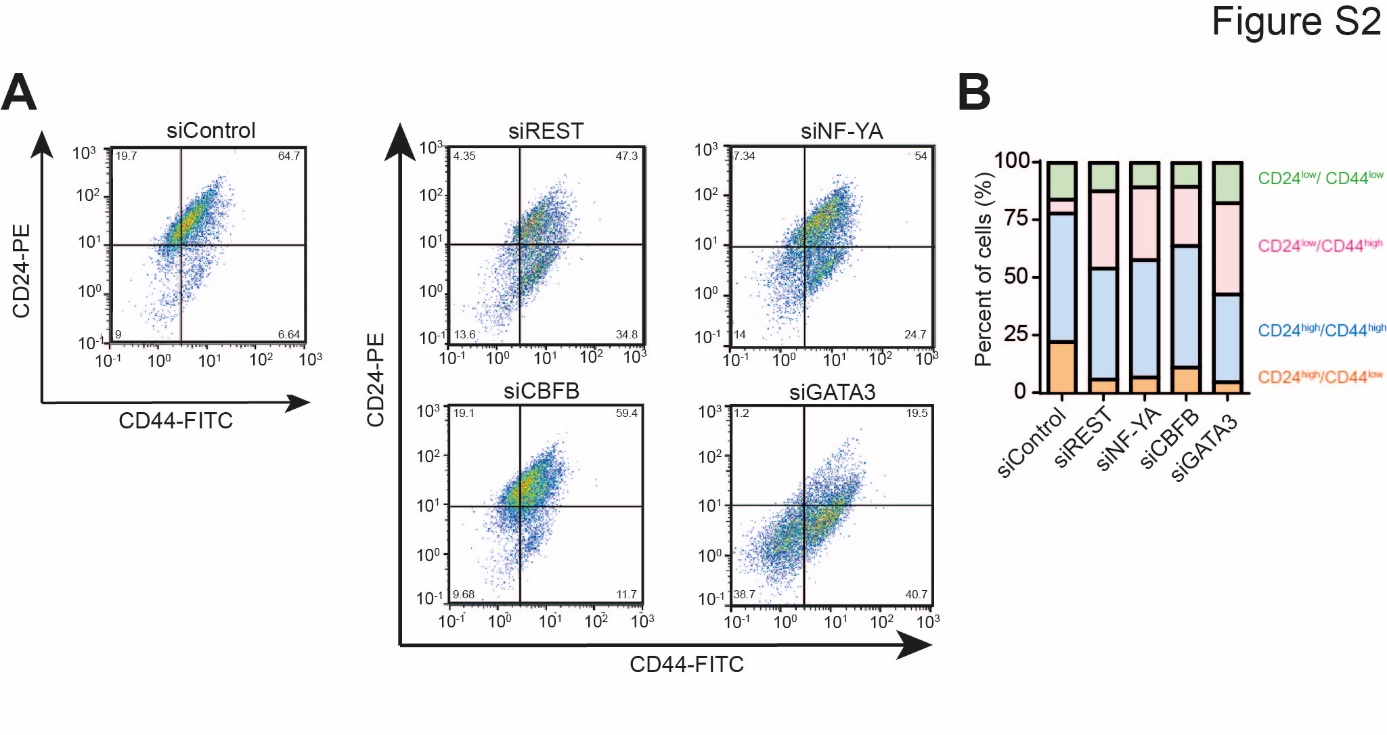

**Figure S3. CD44 and CD24 profiles following siRNA against REST, NF-YA, CBFB and GATA3**

**A,** Representative dot plots are shown from flow cytometry for CD24 and CD44 following transfection of either siControl, or siREST, siNF-YA, siCBFB and siGATA3 as described in Materials and Methods. **B,** Quantification of the different sub-population obtained for each condition.

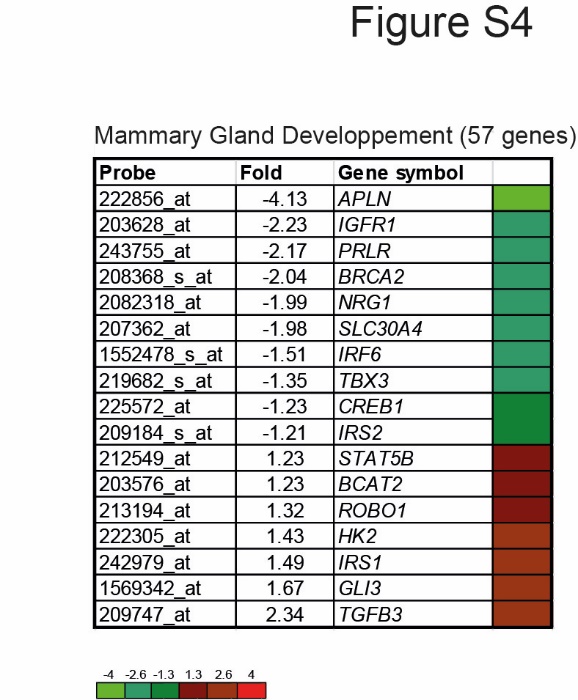

**Figure S4. Genes identified as involved in Mammary Gland development**

Data resulted from the transcriptomic analysis of genes regulated during mammary gland development following miR-106a-3p expression.

**Table S1** List of the primer sequences used in the study.

| **Gene Name** | **Sequences** |
| --- | --- |
| *CBFB*-F | CCCGACCAGAGAAGCAAGTT |
| *CBFB*-R | GCCACAAAAGCGATTTCCGA |
| *NFYA*-F | CAGCAATAGTTCGACAGAGCAGA |
| *NFYA*-R | CTACCTGGAGGGTCTGGACTTG |
| *REST*-F | GGAGGAAACATTTAAGAAACCATTT |
| *REST*-R | CATGGCGGGTTACTTCATGT |
| *GAPDH*-F | CCTGGTATGACAACGAATTT |
| *GAPDH*-R | GTGAGGGTCTCTCTCTTCCT |
| *OCT3/4*-F | TTCAGCCAAACGACCATCTG |
| *OCT3/4*-R | CACGAGGGTTTCTGCTTTGC |
| *SOX2*-F | CCCACCTACAGCATGTCCTACTC |
| *SOX2*-R | TGGAGTGGGAGGAAGAGGTAAC |
| *NANOG*-F | TTCCCTCCTCCATGGATCTG |
| *NANOG*-R | TGTTTCTTGACTGGGACCTTGTC |
| *SMAD2-F* | CCATCGAAAGGATTGCC |
| *SMAD2-R* | ATTCGCAGTTTTCAATTGCC |
| *MMP2-F* | CCCCCAAAACGGACAAAG |
| *MMP2-R* | CCGCATGGTCTCGATGGTAT |
| *ZEB2-F* | AAGCCAGGGACAGATCAGC |
| *ZEB2-R* | CCACACTCTGTGCATTTGAACT |
| *N-CADHERIN-F* | CTCCATGTGCCGGATAGC |
| *N-CADHERIN-R* | CGATTTCACCAGAAGCCTCTAC |
| *SNAIL1-F* | CCCAGTGCCTCGACCACTAT |
| *SNAIL1-R* | GCTGGAAGGTAAACTCTGGATTAGA |
| *TWIST1-F* | GGCATCACTATGGACTTTCTCTATT |
| *TWIST1-R* | GGCCAGTTTGATCCCAGTATT |
| *KRT14-F* | ATGACCTTGGTGCGGATTT |
| *KRT14-R* | ATCGAGGACCTGAGGAACAA |
| *TGFBR2-F* | GCAGGTGGGAACTGCAAGAT |
| *TGFBR2-R* | GAAGGACTCAACATTCTCCAAATTC |
| *TGFB1-F* | GCCGCTGCCCATCGTGTACTA |
| *TGFB1-R* | GGCTTGGGCACGGGTGTCC |
| *MMP9-F* | GAACCAATCTCACCGACAGG |
| *MMP9-R* | GCCACCCGAGTGTAACCATA |
| *ZEB1-F* | GGGAGGAGCAGTGAAAGAGA |
| *ZEB1-R* | TTTCTTGCCCTTCCTTTCTG |
| *VIMENTIN-F* | TACAGGAAGCTGCTGGAAGG |
| *VIMENTIN-R* | ACCAGAGGGAGTGAATCCAG |
| *B-CATENIN-F* | GGGAGACGGAGGAAGGTCTGA |
| *B-CATENIN-R* | CACGCTGGATTTTCAAAACAGTTG |
| *E-CADHERIN-F* | GCCGAGAGCTACACGTTCA |
| *E-CADHERIN-R* | GACCGGTGCAATCTTCAAA |
